# Path integration impairments reveal early cognitive changes in Subjective Cognitive Decline

**DOI:** 10.1101/2025.02.17.638583

**Authors:** Vladislava Segen, Md Rysul Kabir, Adam Streck, Jakub Slavik, Wenzel Glanz, Michaela Butryn, Ehren Newman, Zoran Tiganj, Thomas Wolbers

**Author notes:** Corresponding author: Vladislava Segen. Equal senior authors (Zoran Tiganj, Thomas Wolbers).

## Abstract

Path integration, the ability to track one’s position using self-motion cues, is critically dependent on the grid cell network in the entorhinal cortex, a region vulnerable to early Alzheimer’s disease pathology. In this study, we examined path integration performance in individuals with subjective cognitive decline (SCD), a group at increased risk for Alzheimer’s disease, and healthy controls using an immersive virtual reality task. We developed a Bayesian computational model to decompose path integration errors into distinct components. SCD participants exhibited significantly higher path integration error, primarily driven by increased memory leak, while other modelling-derived error sources, such as velocity gain, sensory and reporting noise, remained comparable across groups. Our findings suggest that path integration deficits, specifically memory leak, may serve as an early marker of neurodegeneration in SCD and highlight the potential of self-motion-based navigation tasks for detecting pre-symptomatic Alzheimer’s disease-related cognitive changes.

**Teaser:** Virtual reality, modelling, and plasma biomarkers reveal path integration deficits in pre-symptomatic Alzheimer’s vs normal aging.

## Introduction

Spatial navigation is a multifaceted behaviour involving various cognitive processes such as memory storage and retrieval, multisensory integration, and decision-making. Central to navigation is path integration (PI), a process of continuously updating one’s position and orientation based on the integration of self-motion cues(*1*). This mechanism is crucial for the development of cognitive maps, aiding in the association of environmental cues with location estimates(*2*). PI is thought to critically depend on grid cell computations in the entorhinal cortex (EC)(*3*), which is also the first neocortical region to exhibit tau pathology and neurodegeneration in Alzheimer’s disease (AD)(*4*). Consistent with these findings, impaired grid cell dynamics and navigation deficits are evident early in AD mouse models(*5, 6*). In humans, young APOE-e4 carriers, a known risk factor for AD, have shown altered grid-like BOLD signals(*7*). Moreover, behavioural work has suggested corrupted PI in patients with Mild Cognitive Impairment (MCI) and early AD(*8, 9*), particularly in cases when AD-related pathology is present. Howett et al.(*9*) even demonstrated that (i) PI performance was more sensitive at discriminating between AD biomarker positive vs. negative MCI patients compared to standard neuropsychological assessments, and (ii) that PI performance was related to CSF tau and EC volume, further outlining the link between PI and AD-related pathology.

Despite the evidence that PI is affected in MCI and early AD, it remains unknown whether PI deficits emerge at earlier stages of the disease, before traditional cognitive symptoms become apparent. Earlier identification is particularly important as it opens a window for potential interventions at a stage when treatment could be more effective, potentially altering the disease trajectory(*10, 11*). Subjective Cognitive Decline (SCD) presents a unique opportunity in this regard, because it is increasingly acknowledged as potentially the earliest stage of AD(*12*).

Older adults with SCD self-report cognitive deficits that are not detectable through standard neuropsychological testing(*13*), and they have shown signs of tau pathology in EC(*14*). To date, however, it is unknown if PI is affected in SCD and, if so, what mechanisms may represent the earliest degradation of PI due to emerging AD pathology.

To achieve a nuanced understanding of the mechanisms that could underlie PI impairments in SCD, we developed a hierarchical Bayesian model that decomposes observed PI errors into distinct components. Our model builds upon previous leaky integrator models(*15–17*) that assume a linear accumulation of errors with time or distance, influenced by the leaking of information from the memory trace. Parameters of the model include memory ‘leak’, velocity gain, additive bias, accumulating noise and reporting noise. By incorporating these parameters, the model accounts for noise, representing random fluctuations, and biases, indicating systematic deviations from the true path, both of which contribute to the overall accuracy of the PI process. Unlike previous models that were based on maximum likelihood, which yield point estimates of parameters, the Bayesian approach estimates full posterior distributions(*18*), allowing for a richer quantification of uncertainty. Additionally, its hierarchical structure enables the simultaneous modelling of individual differences and group-level effects, offering deeper insights into the variability of PI impairments in SCD. By incorporating prior information, the Bayesian framework is also more robust to noisy data.

To determine if and how PI is impaired in preclinical AD, we tested patients with SCD and matched controls on an immersive, self-guided virtual reality-based PI task. We eliminated distal cues and utilised curved paths to more accurately replicate continuous PI observed in animal studies, minimising reliance on non-spatial heuristics and configural strategies commonly associated with triangular paths in human experiments(*19–21*). Additionally, given the reports that early AD may be associated with angular deficits(*22*) we complemented our PI task by including a novel response to assess angular integration, without being confounded by distance encoding as in previous studies (e.g.(*22, 23*), see (*19*) for further discussion). To preview, our results show that individuals with SCD exhibit larger PI errors compared to controls, driven by increased memory leak as revealed by computational modelling.

Importantly, these deficits were not associated with differences in angular integration, movement dynamics, or visual distance estimation, underscoring the specificity of PI impairments in SCD.

## Results

Data were collected from 102 participants, comprising 72 controls and 30 individuals with SCD. No significant differences were observed between the groups in terms of neuropsychological assessments, self-reported navigation abilities, and visuo-spatial working memory (Table 1). The SCD group was slightly older (BF □ □ = 1.916), and controls performed slightly better on the gait assessment (BF □ □ = 3.057), although both groups scored near ceiling (12-point maximum).

**Table 1.** Demographic characteristics

|  | **Control** | **SCD** | **BF10** |
| --- | --- | --- | --- |
|  | **Mean (SD)** | **Mean (SD)** |  |
| Age | 65.5 (5.68) | 68.7 (7.76) | **1.916** |
| MoCA | 27.0 (1.80) | 26.7 (2.00) | 0.274 |
| Self-reported spatial abilities | 69.4 (22.80) | 78.5 (24.8) | 0.868 |
| Visuo-spatial working memory (corsi block task) | 4.5 (0.96) | 4.58 (0.898) | 0.236 |
| Gait (subset of functional gait assessment task(24)) | 11.3 (1.01) | 10.6 (1.40) | **3.057** |
| Completed PI trials | 79.4 (14.70) | 79.2 (18.6) | 0.227 |

To measure PI, participants engaged with an immersive virtual reality environment through a head-mounted display (an example video is available in supplementary materials). They navigated the environment using self-motion cues (vestibular, proprioceptive, motor efference copies and optic flow). For the PI task (Fig. 1), participants followed a floating object along eight distinct pre-defined curved paths (Fig. S1). They were required to report two key metrics at designated stopping points (Stop 1 and Stop 2, Fig. 1): 1) initial heading orientation (angular integration [AI] response), and 2) distance and direction back to the start of the path (PI response). Some trials featured only a single stopping point at the end of the path (Fig. 1; see Methods). After outlier exclusion, both groups presented a comparable number of valid trials for analysis (Table 1).

**Fig. 1.**
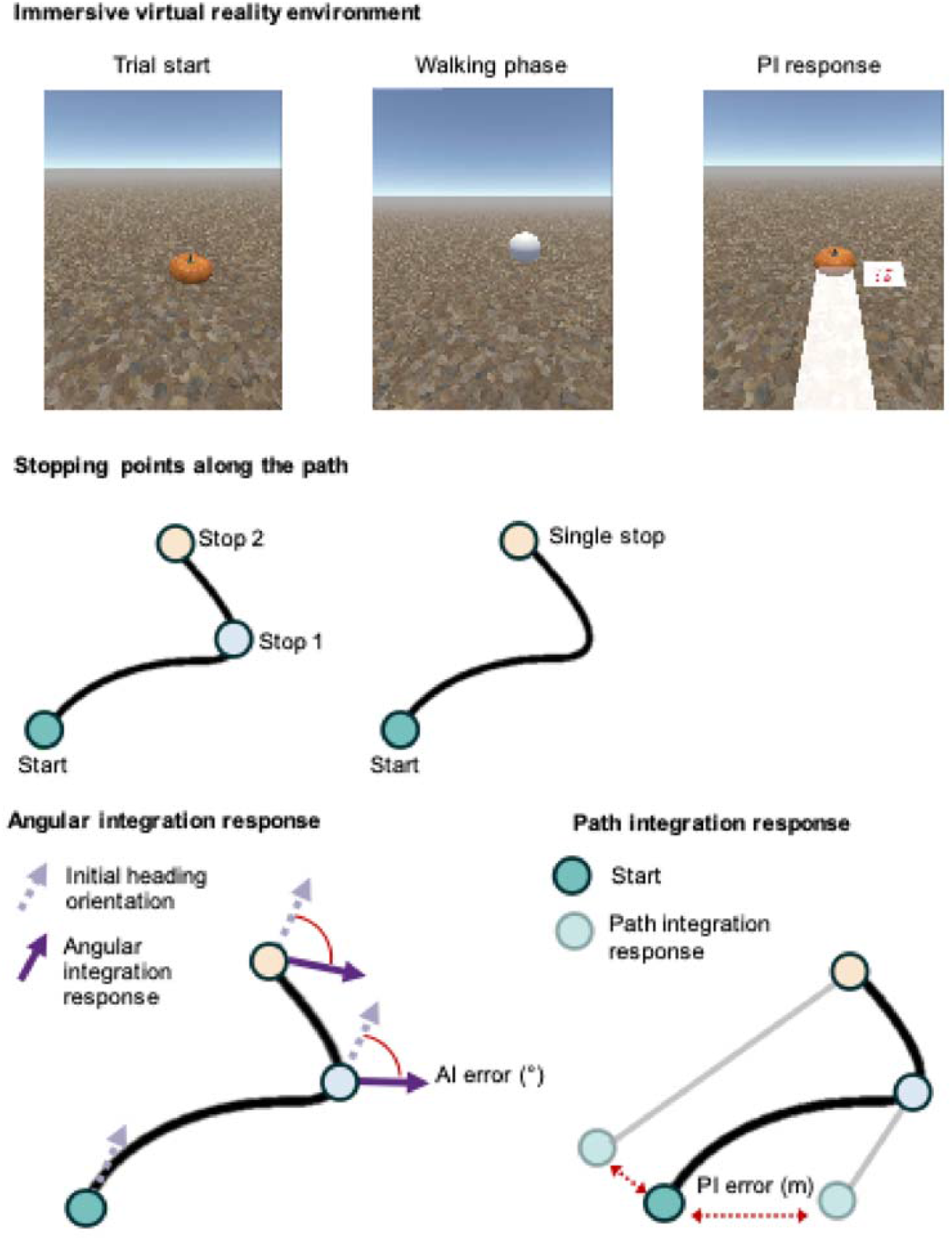
Task Schematic for Path Integration and Angular Integration. (Top) **Example of the immersive virtual reality environment illustrating the key stages of the task.** Participants started at a designated point marked by a visible object (e.g., a pumpkin). They then followed a curved path by walking towards a floating sphere (the object was no longer visible). At the stopping point, they performed the path integration response by repositioning the object to its original location. This was done by turning and estimating the distance using a white line displayed on the ground within the virtual environment. Participants also saw a numerical representation of the response line length. Additionally, participants performed an angular integration response (not shown) by rotating to their initial heading orientation (see bottom panel). (Middle) **Example of a curved path**, performed either with two stopping points, Stop 1 in the middle of the path and Stop 2 at the end (left), or with a single stop at the end of the path (right). (Bottom) **Representation of key task elements and metrics**. (Left) Participants angular integration (AI) response example, where participants are asked to indicate their initial heading orientation at each stopping point by rotating their head and body. This task required participants to memorize their initial heading and update that heading as they navigate the curved path. Dashed arrow represents the initial heading orientation and solid purple arrow represents the AI response. The absolute difference between the two represents AI error. (Right) For the path integration (PI) response, participants were asked to indicate the start position of the path by turning to the “presumed” start location and then indicating distance to start. The difference between the start location and the PI response indicates PI error (m). Participants performed both responses (angular and path integration) at each stopping point.

### SCD patients show reduced PI performance

Using a regression model, for group and stopping point with age, sex and MoCA scores as covariates, we found that participants with SCD exhibited larger PI errors compared to healthy controls (Estimate =0.257 SE = 0.065, t =3.925, p<0.001, Fig. 2A). Both groups demonstrated higher PI error at the 2nd stopping point at the end of the path relative to the intermediate response points (Estimate = 0.560, SE = 0.090, t = 6.245, p<0.001, Fig. 2B). Critically, there were no significant differences in PI errors for the final stop between trials with or without intermediate stopping points for either group (t=1.238, p=0.217), suggesting that in both groups, errors increased with increasing walked distance from the start location. Replicating previous findings, PI errors increased with advancing age (Estimate = 0.427, SE = 0.063, t = 6.728, p<0.001, Fig. 2D), and females exhibited higher PI errors than males (Estimate = 0.306, SE = 0.067, t = 4.557, p<0.001, Fig. 2C). Full regression results are reported in supplementary materials (Table S1). We assessed whether participants performed better than chance on the PI task. Both groups outperformed chance at the first stopping point. Importantly, at the final stopping point, SCD participants did not perform above chance, while controls maintained above-chance performance in both trials with and without the intermediate stopping point (Fig. S2).

**Fig. 2.**
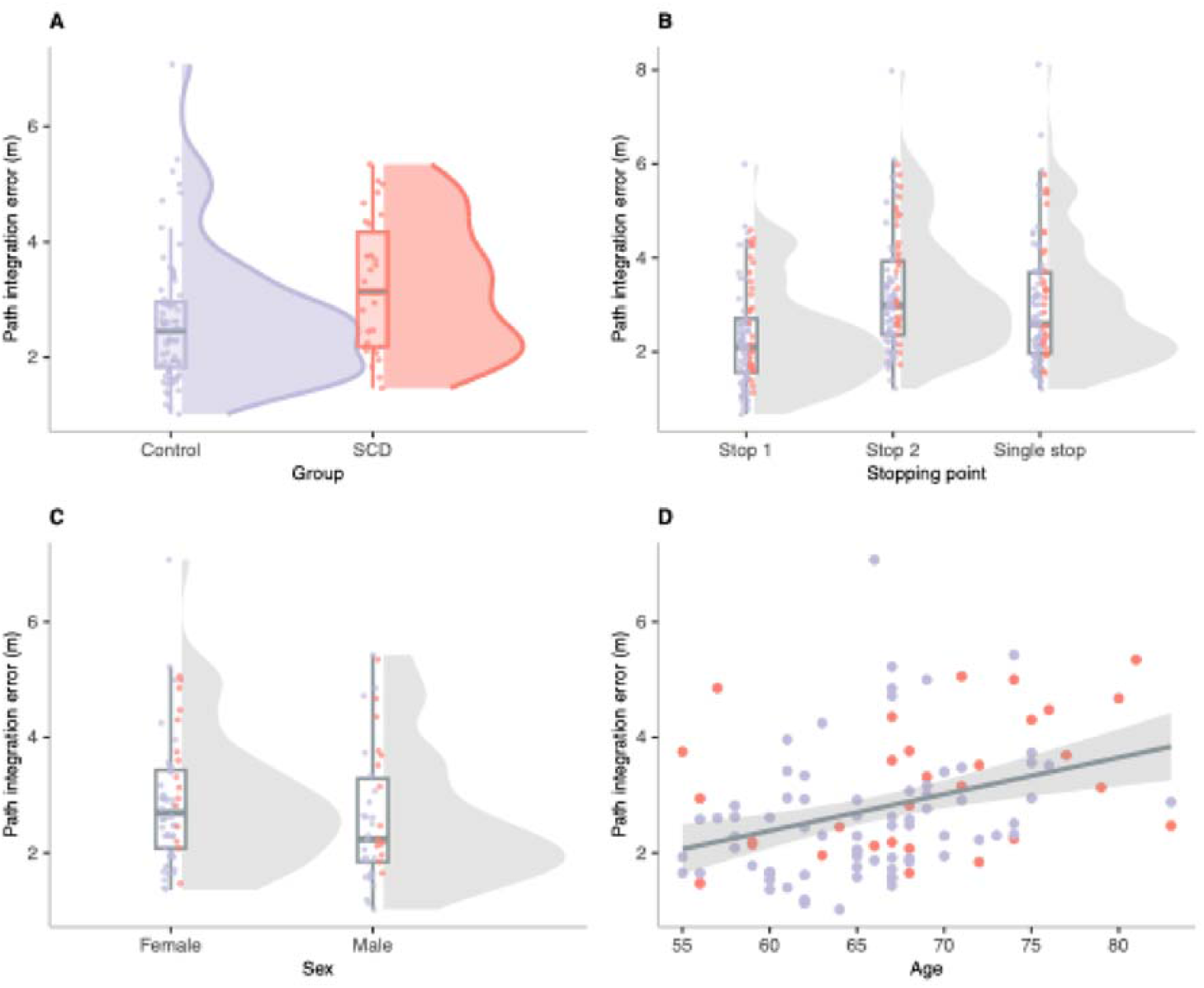
Path integration performance. **A**) Group differences in PI error; healthy controls exhibited significantly lower PI errors compared to the SCD group, **B**) with errors increasing at the final stopping point relative to intermediate points in both groups. **C**) Sex differences in PI error; females exhibited significantly higher PI errors compared males. Boxplots show the median and interquartile range (IQR); violins indicate data distribution with individual data points **D**) PI error increased as a function of age across both groups, the shaded area represents the 95% confidence interval of the regression line. All plots are based on robust multiple linear regression models (p<0.05).

In contrast to PI error, there were no differences in AI error between the groups (BF□□ =0.270, Fig. 3A), with both groups performing significantly better than chance (Control: mean= 46.126°, BF□□ = 50.125; SCD: mean= 49.463°, BF□□ = 16.043). Similar to PI error, we found higher AI error between the 2nd stopping point at the end of the path relative to the intermediate response points (Estimate = 11.354°, SE = 1.881, t = 6.04, p<0.001, Fig. 3B).

**Fig. 3.**
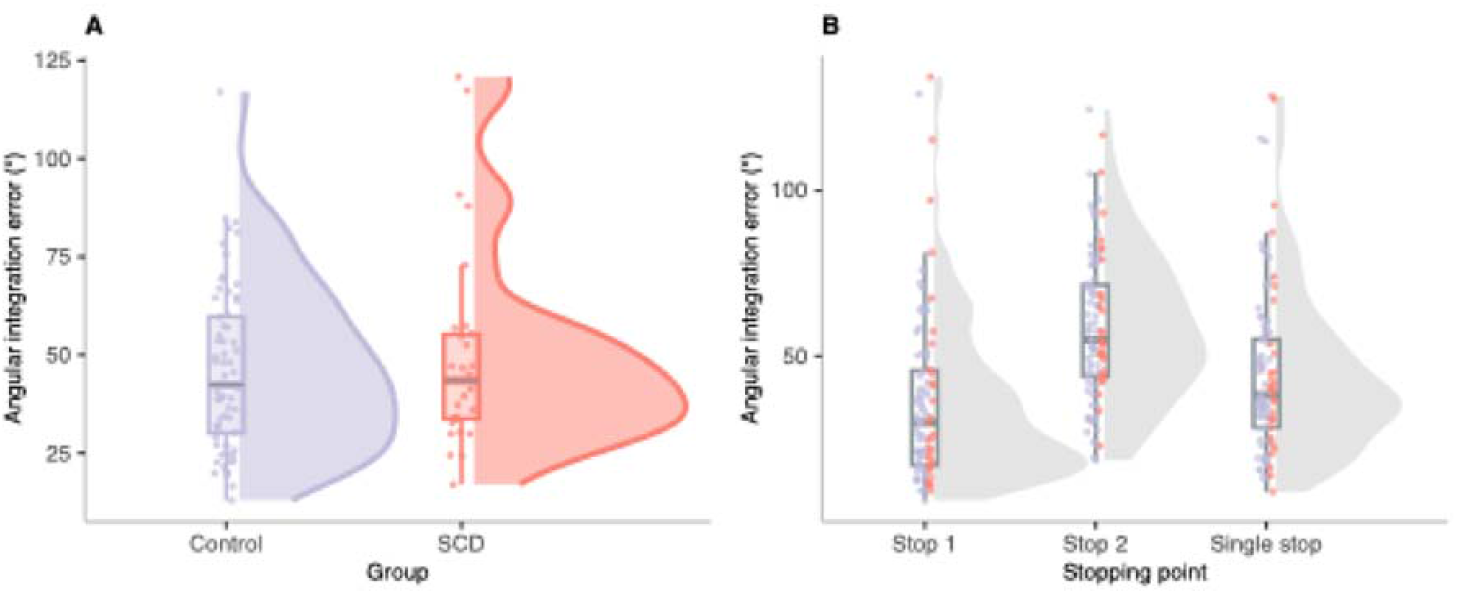
Angular integration performance. **A**) No significant group differences were observed in angular integration (AI) error between healthy controls and individuals with SCD. **B**) AI error varied across stopping points, with higher error at Stop 2 compared to Stop 1, and lower error at the final stopping point in trials with only a single stop compared to those with an intermediate stop. Boxplots show the median and interquartile range (IQR); violins indicate data distribution with individual data points. All plots are based on robust multiple linear regression model.

Finally, AI error was associated with increasing age and was higher in females compared to males (Table S2), but there was no link between AI error and corsi block span – a measure of visuo-spatial working memory (Figure S3).

### Movement characteristics and visual distance perception are unlikely to drive PI differences between groups

To test whether group differences in PI error were driven by movement dynamics, we compared head movements, angular and translational velocity, and head pitch (Fig. S4) using Bayesian t-tests, assessing evidence for the null hypothesis. Controls and SCD neither differed in head movements during walking, (BF□□=4.049) nor in translational (BF□□=4.129) and angular velocities (BF□□=2.035).

We further examined whether SCD participants sampled the environment differently by looking downward more frequently during walking, which could impair optic flow perception(*25, 26*). Since gaze behaviour was not recorded, head pitch data from the HMD served as a proxy, revealing no group differences (BF□□=2.034). Together, with all analyses yielding BF□□ >1, we conclude that movement dynamics are unlikely to contribute to the differences in PI error between groups. These findings are consistent with previous work by Stangl et al. (17), who also reported no differences in movement dynamics between young and older adults, supporting the notion that PI errors in aging and early AD are not primarily driven by altered sampling of the environment but rather by underlying changes in spatial computation.

Next, we examined changes in PI performance and movement dynamics from early to late trials (comparing the first 10% vs. last 10% of trials; Fig. S5) to ensure no differences in learning dynamics or task adaptation between groups, which could confound PI performance interpretations. First, we did not find any changes in PI performance between early and late trials (estimate = 0.006, SE = 0.093, t = 0.068, p = 0.946), with no significant interaction between group and trial stage (estimate =-0.046, SE = 0.093, t = -0.493, p = 0.623). In terms of movement dynamics, we observed an increase in translational and angular velocity between early and late trials (Translation: estimate = 0.026, SE = 0.003, t = 10.033, p < 0.001; Angular: estimate = 1.807, SE = 0.243, t = 7.452, p < 0.001), with similar patterns for path groups (Translation: p=0.840; Angular: p= 0.983). Additionally, both groups showed an overall decrease in head movements in later trials (estimate = -96.101, SE = 14.688, t = -6.543, p < 0.001), potentially reflecting that participants realised the futility of extensive head movements due to the lack of distal cues in the environment, with no interaction between group and trial stage (p=0.210). Finally, head pitch remained unchanged across trials (p= 0.781), with no group or trial stage interactions (p=0.639).

In addition to the PI task, we included a distance estimation task to assess potential differences in visual distance perception and response precision between control and SCD participants.

Participants memorised and reproduced distances to an object (1.4, 3.4, and 5.9 metres) using a virtual ruler. We found no significant group differences in distance estimation (estimate = 0.027, SE = 0.024, t = 1.113, p = 0.267, Fig. S6), suggesting comparable visual distance perception and estimation across groups. Both groups exhibited a Weber’s law-like effect, with error increasing as the distance increased from 1.4 to 3.4 metres (estimate = 0.288, SE = 0.033, t = 8.722, p < 0.001) and further from 3.4 to 5.9 metres (estimate = 0.351, SE = 0.033, t = 10.553, p < 0.001). Notably, the SCD group exhibited a larger increase in error at the longest distance compared to controls (estimate = 0.103, SE = 0.033, t = 3.092, p < 0.001), suggesting they may experience more pronounced Weber-like uncertainty at greater distances. However, comparison of the beta estimates indicated that the main effect of distance (β = 0.351) was more than three times larger than the group × distance interaction (β = 0.103), suggesting that increasing distance impacted both groups more strongly than the group difference alone.

Additionally, distance error increased with age (estimate = 0.007, SE = 0.003, t = 1.978, p = 0.049), although this effect was borderline significant and should be interpreted with caution. Full results are reported in Table S3.

### Characterising error sources with a computational model

To better understand the mechanisms that contribute to the observed PI errors, we developed an extended computational model based on the distance-based framework introduced by Stangl et al.(*17*). This enhanced model addresses gaps in prior approaches by capturing both individual variability and shared characteristics of healthy aging and early pathological changes (i.e., SCD). Our model simulates participants’ internal location estimates during PI using a two-dimensional diffusion equation, incorporating, velocity gain(*α*), memory leak (*β*), additive bias (**b**) and accumulating noise (*σ*_0_). Internal estimates are generated based on reported distance 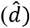 and angle, with addition of Weber-like reporting noise (*σ* _r_)drawn from a normal distribution with zero mean and standard deviation proportional to the reported distance 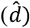.

To infer the model parameters ( *α,β*, **b**, *σ*_O_, *σ*_r_), we utilized a Bayesian hierarchical approach, which provides distinct advantages over traditional methods based on likelihood maximization. Specifically, this approach accounts for individual variability while capturing shared group-level characteristics. The Bayesian framework allows for prior knowledge integration and robust parameter estimation via posterior distributions. Parameter inference was conducted using Markov Chain Monte Carlo (MCMC) sampling with the No-U-Turn Sampler (NUTS), ensuring efficient exploration of the parameter space and reliable posterior estimates(*27*). This model effectively captures variability across individuals and groups, enhancing our understanding of cognitive changes in aging and SCD.

### Model selection/evaluation

To determine the most parsimonious model, we compared candidate models combining various error sources (Fig. 4). Model complexity and fit were assessed using expected log predictive density for leave-one-out cross-validation (elpd_loo_)(*28*). The full model yielded the highest elpd_loo_ value, indicating the best numerical fit, and was therefore retained as the primary model for explaining PI error sources. However, we note that several reduced models—specifically those omitting additive bias, accumulating noise, or both—had elpd_loo_ values that were similar to the full model. To ensure our findings were not dependent on model choice, we re-ran all group-level comparisons using these alternative models. The results remained consistent, supporting the robustness of our conclusions. These additional analyses are reported in the supplementary materials (Figure S7).

**Fig. 4.**
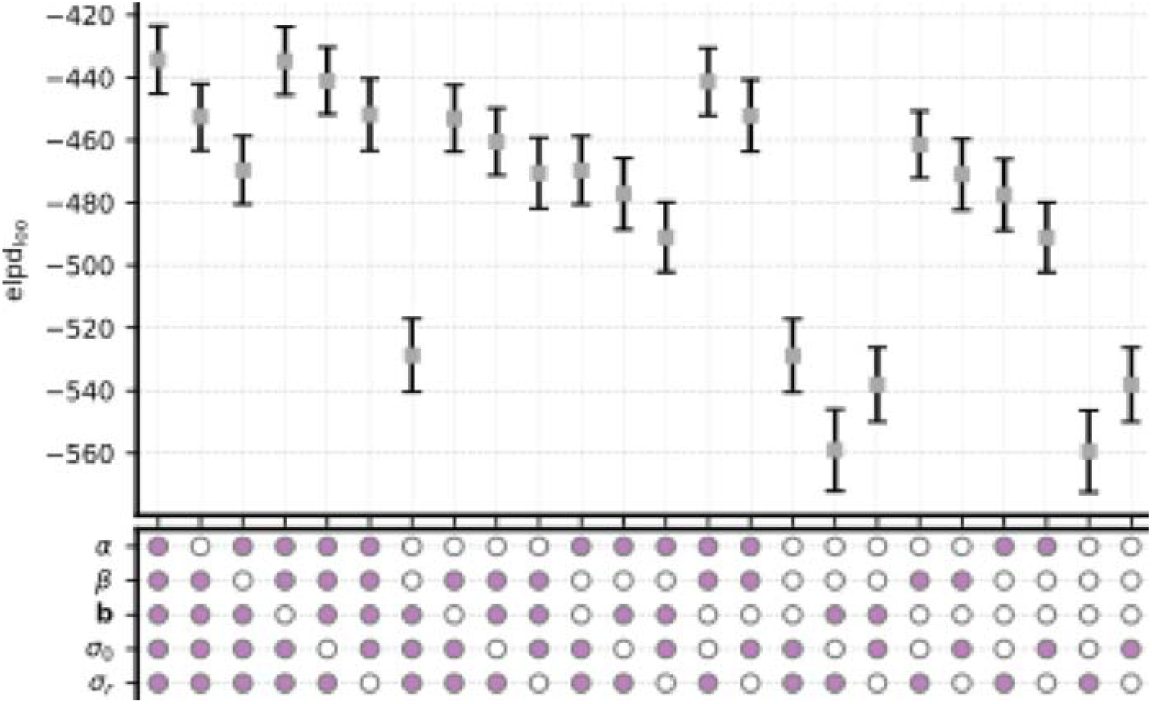
Comparison of Candidate Models Across Error Sources. Comparison of candidate models incorporating different combinations of error sources: velocity gain (α), memory leak (β), additive bias (), accumulating noise (), and reporting noise (). Error sources included in each model are represented below the graph as filled (purple). The expected log pointwise predictive density for leave-one-out cross-validation (elpd_loo_) is shown for each model (mean ± SEM). Models with higher elpd_loo_ values indicate better predictive performance. The “full” model demonstrates the best fit to the data (highest elpd_loo_ value).

### Memory leak distinguishes SCD patients from healthy controls

What are the mechanisms that may have caused increased PI errors in individuals with SCD? To address this question, we first calculated mean parameter estimates for each participant and compared them using linear regression with age and group as covariates (results reported in Table S4). We found that SCD participants exhibited significantly higher memory leak than Controls (β; estimate = 0.055, SE = 0.021, t=2.664, p=0.009, Fig. 5B), indicating a greater tendency for stored information to decay over travelled distance. We also observed a marginally significant group difference in reporting noise (σ*_r_^2^*; estimate = 0.035, SE = 0.017, t = 2.070, p = 0.042, Fig. 5E), with SCD participants exhibiting slightly higher values compared to controls. In contrast, there was no evidence of a significant group difference in velocity gain (α; estimate = -0.026, SE = 0.051, t = -0.518, p = 0.606, Fig. 5A), additive bias (||**b**||; estimate = 0.001, SE = 0.003, t = 0.570, p = 0.571, Fig. 5C), and accumulating noise (σ*_o_^2^*; estimate = 0.016, SE = 0.025, t = 0.646, p = 0.520, Fig. 5D). Across both groups, age was associated with increases in memory leak (β; estimate = 0.008, SE = 0.003, t = 2.765, p = 0.007, Fig. S8A) and reporting noise (σ*_r_^2^*; estimate = 0.008, SE = 0.002, t = 3.361, p = 0.001, Fig. S8B).

**Fig. 5.**
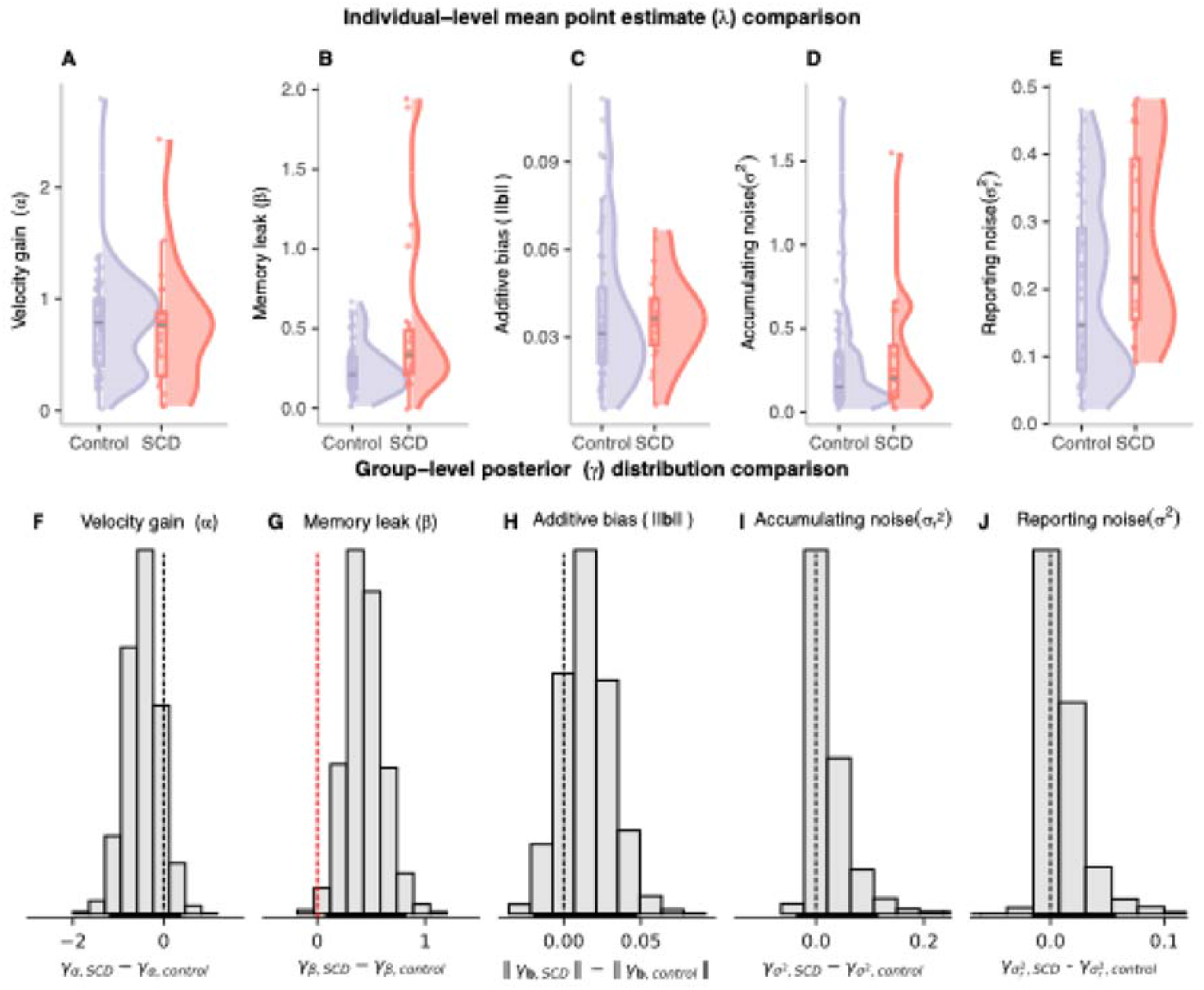
Comparison of Computational Model Parameters Between Control and SCD Groups. Upper panel (**A-E**) Violin plots show the distribution of individual-level mean point estimates, overlaid with boxplots indicating the median and interquartile range (IQR) and individual data points for velocity gain (α), memory leak (β), additive bias (‖**b**‖), accumulating noise σ *^2^*), and reporting noise (σ *^2^*) for Control (blue) and SCD (red) participants. Compared to Controls, SCD participants exhibited significantly higher memory leak (β) (**B**) as well as marginally higher reporting noise (σ*_r_^2^*) (**E**) while other parameters (α, ‖**b**‖, σ*_0_^2^*) showed no significant group differences. Asterisk indicates significant effect (** p <0.01, * p< 0.05) from a robust linear regression model with age as a covariate. Lower panel (**F-J**) Posterior distributions of the differences between control and SCD groups for the group-level mean parameter γ. The horizontal bars near the x-axis denote the 95% Highest Density Interval (HDI) of the posterior distributions for group differences. Dashed vertical lines indicate zero, and the percentages reflect the proportion of the posterior distribution on either side of zero, providing evidence for the likely direction of group differences. (**G)** The posterior distributions revealed strong evidence for higher memory leak (β) in individuals with SCD compared to Controls (red dashed line), with 99.7% of the distribution above zero and a 95% HDI excluding zero, indicating a statistically credible group difference. In contrast, posterior distributions of γ for the remaining parameters—velocity gain, additive bias, accumulating noise, and reporting noise (**F, H-J**)—showed negligible evidence for group differences, as their 95% HDIs overlapped zero (black dashed line).

To further assess the robustness of our findings, we examined group differences in PI error sources using the Highest Density Intervals (HDIs) of the posterior distributions of the group-level mean model parameters (see Fig. S9). HDIs provide a comprehensive summary of parameter differences by capturing the most credible range rather than relying solely on point estimates, offering a clearer representation of uncertainty and group differences. Consistent with the individual-level analysis, the differences in the posterior distributions of for memory leak (*β*) provide strong evidence for higher values in individuals with SCD compared to controls, with 99.5% of the distribution above zero (Fig. 5G). In addition, the 95% HDI [0.10,0.79] did not include zero, suggesting a statistically credible and significant group difference. While we observed a marginally significant group difference in reporting noise (σ_r_^2^) at the individual level (p = 0.042), this was not supported at the group level, where the posterior distribution overlapped with zero (Fig. 5J). This suggests that the group difference in reporting noise may be less robust and should be interpreted with caution. No evidence for group differences was found for the remaining parameters, because the 95% HDIs for velocity gain (*γ_a_*), additive bias ( *γ_b_*), accumulating noise 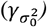 (Fig. 5F, 5H, 5I) all overlapped zero. A subsequent ROPE analysis ([-0.1, 0.1]) supported practical equivalence for the remaining parameters, as most of the 95% HDI samples fell within these bounds(*29*). Together, these findings highlight **memory leak** as the most consistent and reliable parameter distinguishing individuals with SCD from healthy controls, whereas other error sources—such as reporting noise—were either less robust or comparable across groups.

To ensure that our findings were not driven by prior assumptions, we tested a range of reasonable hyperparameter values for the memory leak parameter (β). Specifically, we re-ran the hierarchical model using different priors and found that the group-level difference between SCD and control participants remained consistent. Full results of this sensitivity analysis are provided in the supplementary materials (Figure S10).

### Blood NFL Predicts PI Errors, Velocity Gain Deviations, and Increased Reporting Noise

We also obtained plasma-based biological biomarker data related to neurodegeneration from a subset of participants (SCD=27, Control=54). Specifically, we measured plasma levels of neurofilament light chain (NFL), a marker of general neurodegeneration(*30, 31*), and pTau181, associated with AD-related tau accumulation(*31, 32*). We also included APOE (ε4 carriers and ε4 noncarriers), a risk factor for AD(*33*), in the analysis. Our analysis of these plasma biomarkers showed no significant differences in the concentrations of NFL (BF[[=0.461, Fig. 6B) as well as no differences in the number of ε4 carriers and ε4 noncarriers between Control and SCD groups (χ ^2^p=0.796, Fig. 6C).

**Fig. 6.**
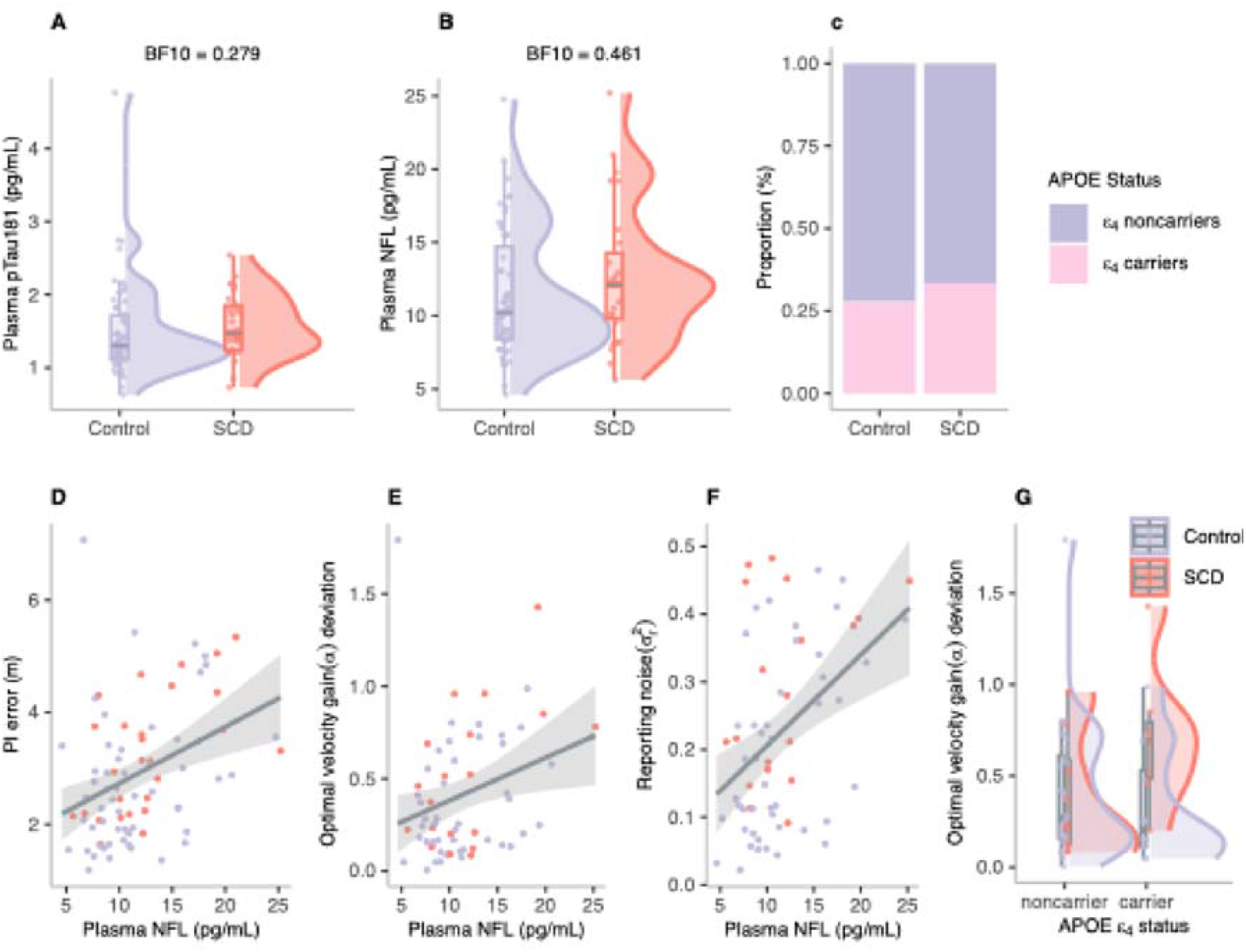
Plasma Biomarkers, APOE Genotype, and Associations with PI and Error Sources. (**A-B**) Violin plots showing plasma levels of pTau181 and NFL in Control (blue) and SCD (red) groups. Bayesian t test analyses provided evidence supporting no group differences in pTau181 (BF□□ =0.279) and NFL (BF□□ =0.461). (**C**) Proportion of APOE ε4 carriers (pink) and noncarriers (blue) across Control and SCD groups, showing no significant differences (p=0.796) based on the chi squared test. (**D–F**) Scatter plots illustrating the predictive relationship between plasma NFL levels and behavioural outcomes. Shaded areas represent the 95% confidence interval for regression lines. Higher NFL levels were associated with increased PI error (**D**), greater deviations from the optimal velocity gain (**E**), and higher reporting noise (**F**). (**G**) Significant interaction between APOE ε4 status and group in predicting deviation from optimal velocity gain, with greater deviations observed in SCD ε4 carriers. Boxplots show the median and interquartile range (IQR); violins indicate data distribution with individual data points. Plots **D-G** are based on robust multiple linear regression models.

Next, we investigated the predictive relationship between PI error and blood-based biomarkers, with age included as a covariate. While no group differences in biomarker concentrations were observed, biomarkers may still have differential effects across groups, therefore, we included group as an interaction term in the model. NFL was the only significant predictor of increased PI error (Fig. 6D; estimate = 1.080, SE = 0.366, t = 2.954, p =0.004). Subsequently, to understand the potential biological underpinnings driving distinct error sources contributing to impaired PI we examined if these biomarkers predict individual parameter estimates derived from the computational model, and whether there are differential effects across groups. Higher NFL levels were significantly associated with greater deviations from optimal velocity gain (Fig. 6E; absolute deviation from α = 1; estimate = 0.138, SE = 0.042, t = 3.277, p = 0.002).

We also observed a significant interaction between group and APOE ε4 status in predicting velocity gain deviation, such that SCD participants who were APOE ε4 carriers showed larger deviations (estimate = 0.094, SE = 0.044, t = 2.113, p = 0.039, Fig 6F). However, this result should be interpreted with caution due to the limited number of SCD participants with biomarker data (n = 19), and especially APOE ε4 carriers (n = 7). In addition, NFL levels were also predictive of increased reporting noise (Fig. 6G; estimate = 0.053, SE = 0.019, t = 2.869, p = 0.006). No other biomarkers significantly predicted PI error sources. Full results of PI error and modelling parameters analysis in relation to blood biomarkers are reported in Table S5 and S6.

## Discussion

In this study, we examined path integration in individuals with SCD and healthy controls using a self-guided immersive virtual reality task. SCD participants showed significantly higher PI errors than controls. A hierarchical Bayesian model revealed that these deficits were primarily driven by increased memory leak, with a marginally higher reporting noise in SCD, while other parameters—velocity gain, additive bias, and accumulating noise—remained similar between groups. Although no group differences were found in blood biomarkers, NFL, a marker of neurodegeneration, was significantly associated with increased PI errors, velocity gain deviations, and reporting noise.

To the best of our knowledge, this study provides the first evidence for PI impairments in SCD participants, despite their comparable performance to healthy controls on the AI component of the task and in other cognitive domains. Bayesian analyses did not reveal any group differences in head movements, translational and angular velocity, or head pitch, indicating that PI deficits were unlikely to be driven by variations in movement dynamics or sampling strategies, such as a tendency to look downward during navigation. Additionally, both groups exhibited similar changes in performance and movement metrics from early to late trials, with no evidence of group differences in learning or task adaptation. Thereby, our results highlight that PI may uniquely tap into subtle changes in neural computations that are difficult to detect with standard cognitive measures, highlighting its potential as a sensitive marker of pre-symptomatic AD.

It is important to note that our experimental design was specifically tailored to reduce potential confounds often seen in PI tasks. By requiring participants to rely primarily on multisensory self-motion cues (vision, proprioception, vestibular feedback and motor efference copies), we minimized the influence of sensory degradation, which is commonly observed with aging and can impair performance when limited sensory modalities are available(*23, 34, 35*).

Furthermore, the task excluded proximal and distal landmarks(*7, 9, 36*), reducing the likelihood of compensatory landmark-based navigation or reliance on non-spatial heuristics. These design choices create a more “pure” PI task, where older adults had to continuously update their position in space relying on idiothetic cues. The observed deficits in SCD participants, therefore, likely reflect genuine impairments in PI rather than alternative cognitive or sensory alterations.

To gain a deeper understanding of the mechanisms contributing to the overall PI deficits, we developed a hierarchical Bayesian model that decomposes observed PI errors into distinct components. Critically, we found that **memory leak** was the only parameter that reliably distinguished older adults with SCD from healthy controls. Memory leak, as defined in our model, refers to the gradual decay of the state variable, specifically the homing vector encoding the distance and direction back to the starting point as distance increases during path traversal. Our behavioural findings support that this decay occurs over space rather than time, as indicated by the comparable PI performance at the end of the path in trials with and without intermediate stopping points. Notably, trials without intermediate stops had similar distances but shorter durations, emphasizing that memory leak is more closely tied to movement itself— emerging when positional changes occur—rather than during stationary periods. Thus, we conclude that memory leak is unlikely to be driven by working memory deficits. This interpretation is further supported by the absence of group differences on the Corsi block task, a standard measure of visuo-spatial working memory(*37*).

Our interpretation of spatially-dependent error accumulation is further supported by findings from Stangl et al. (*17*), who observed similar patterns of PI error across trials with varying numbers of stopping points. In their study, both younger and older adults showed consistent error accumulation as a function of distance travelled, independent of time, as in our task.

These results suggest that spatial decay is a robust and general feature of PI across age groups, reinforcing the idea that the memory leak observed in our SCD participants reflects a disruption of core navigational computations rather than age-related differences in task engagement or cognitive strategy.

We propose that these PI deficits are related to impaired grid cell function, which may be among the earliest functional changes during Alzheimer’s disease progression (*6, 38, 39*). Grid cells serve as a neural integrator for spatial information supporting PI(*3*), and functional changes in this network may impair the brain’s ability to maintain a stable representation of self-location over the course of movement. Animal models of AD show profound loss of grid tuning(*6, 40, 41*). The additional burden of tau pathology in the EC may disrupt the grid cell network’s capacity to prevent “leakage,” amplifying memory decay and making it a key distinguishing feature from healthy aging, where some degree of leak may also be present but to a lesser extent (c.f. higher leak with age as seen in Stangl et al., (*17*)). Consistent with this hypothesis, young APOE ε4 carriers exhibit reduced grid-cell-like tuning(*39*).

While the precise mechanisms as to how AD pathology may lead to greater memory leak remain speculative, we propose several plausible explanations. One possible mechanistic example of how AD pathology could disrupt spatial computations involves altered attractor dynamics within the hippocampal–entorhinal circuit. Grid cell models based on continuous attractor networks create stable spatial maps by maintaining coherent activity patterns representing the organism’s location(*42*). In these networks, each new location estimate relies on the previously encoded spatial state and velocity updates. The stability of these attractor networks could be compromised by AD pathology, which effectively reduces the network’s “energy well,” making attractor states more prone to drift. In such a weakened network, any slight perturbation (e.g., from sensory noise or normal fluctuations in neural firing) can push the representation away from its stable configuration, causing the previously encoded spatial state to degrade more quickly and amplifying PI errors. This instability could be further exacerbated by AD related dysfunction in parvalbumin interneurons, which compromises the inhibitory control needed for precise network dynamics and grid tunning(*43, 44*). Furthermore, the disruption of axonal transport and synaptic function likely contributes to this weakened network state(*45*). Consequently, updating spatial position becomes increasingly difficult, with the internal representation eroding faster than under normal conditions.

An additional mechanism involves disrupted temporal precision in the sequential updating of the PI signal. Accurate tracking of position relies on rhythmic oscillatory processes— particularly theta and gamma bands—to coordinate neuronal ensembles in the entorhinal-hippocampal circuit(*46–52*). AD-related changes in the EC may reduce synchrony between grid cells and head-direction cells or attenuate the amplitude of key oscillations, potentially by disrupting the function of interneurons that regulate these rhythms(*53, 54*). For example, disease related reduction in cholinergic transmission(*55*) disrupt theta-gamma interactions and grid tunning(*56, 57*). Without precisely coordinated neuronal firing, the system may struggle to integrate velocity and orientation cues at the correct moments, thereby compounding small discrepancies over successive steps. This disruption of temporal precision could further destabilize the state variable, contributing to the “leak” observed in SCD. Since PI relies on cumulative updates, even minor disruptions in the running position estimate can have a cascading effect, resulting in progressive loss of spatial information manifesting as a gradual “leak” in spatial memory.

While our findings and mechanistic interpretations are grounded in the domain of spatial navigation, we acknowledge that accumulator-like processes are not unique to PI. Grid cells and the wider entorhinal and hippocampal circuit, although classically associated with spatial coding, have also been implicated in the integration of temporal sequences (e.g., Kraus et al.), and conceptual as well as semantic information (*59–62*). This raises the possibility that similar “leak”-like effects may arise in non-spatial domains. In addition, we recognize that early AD pathology, particularly tau accumulation, may impact a range of integrative cell types beyond grid cells. Future work should therefore examine whether a more general accumulator dysfunction contributes to cognitive decline in SCD, which would support a broader role of entorhinal and hippocampal circuits in early Alzheimer’s-related changes across multiple cognitive domains.

Beyond differences in memory leak, we also observed a marginal group difference in reporting noise, which may point to a more domain-general source of increased uncertainty in SCD. Interestingly, this effect aligns with the group and distance interaction observed in the distance estimation task, where SCD participants showed disproportionately higher errors at the longest distance. While both effects were modest, they may reflect a shared underlying mechanism— such as increased imprecision in encoding or reproducing continuous magnitudes under higher-demand conditions. This pattern is consistent with Weber-like effects, in which estimation error scales with the magnitude of the stimulus. Although not central to our primary findings, these effects suggest that continuous spatial estimates may become noisier in SCD, particularly under conditions when larger distances need to be estimated.

Recent studies have argued that corrupted angular integration maybe a primary driver of early AD-related deficits(*22*) because pre-clinical or prodromal AD (i.e., MCI, APOE4 status, and other AD-related risks) was associated with higher angular errors(*7, 36, 63*). In contrast to these findings, we did not observe group differences in AI between healthy older adults and individuals with SCD. Moreover, both groups performed significantly better than chance on our AI tasks, despite showing clear differences in PI. This discrepancy may be explained by methodological differences in how AI is assessed. Traditional PI tasks, such as triangle completion, infer deficits from distance and angular errors of the homing response, with distance error as the deviation from the actual start point and angular error as the difference between the correct and reported heading. However, mis-encoding of travelled distance during the outbound path can also induce angular error, potentially confounding the interpretation of angular deficits(*19–21*). To address this, we incorporated an additional task in which participants were asked to remember and recreate their initial heading orientation at each response point, allowing us to disentangle angular integration from distance encoding and the combined processes required for PI. Critically, while the task includes a working memory component—requiring participants to remember their initial heading—it also demands continuous updating of orientation as they navigate the path, integrating rotational changes over time. This dynamic process is central to angular path integration and engages mechanisms beyond static memory recall. Supporting this interpretation, we found no significant relationship between AI error and visuo-spatial working memory performance (as measured by the Corsi block span), either in the full sample or within each group.

Our findings of intact AI alongside PI deficits in SCD align with research on AD rodent models. These studies suggest that head direction (HD) cell coding, a critical component for orientation inputs to grid cells(*64*), is preserved for longer than grid cell integrity during the progression of AD(*5, 6*). It is possible that impaired AI becomes more prominent at later stages of disease progression, such as MCI, as supported by recent modelling studies in humans(*22*). Notably, Ying et al.(*5*) demonstrated that although HD cells maintain normal firing properties and tuning curves in AD mice, early-stage AD is characterized by reduced synchrony between HD and grid cells. This suggests that impaired integration of orientation and distance information may underlie early PI deficits, as evidenced by intact AI but disrupted PI in SCD, consistent with the interpretation that the EC is responsible for integrating these inputs.

Contrary to previous research(*7, 36, 63*), we did not observe larger PI deficits in APOE ε4 carriers, a known risk factor for sporadic AD, despite employing PI tasks without orientation cues, which are considered highly sensitive to PI impairments in this group (e.g., Colmant et al.,(*63*)). This discrepancy may be partly explained by complex interactions between APOE status, lifestyle factors, and sex, as suggested by prior studies(*36*). This aligns with evidence that APOE ε4 expression is modulated by various epigenetic(*65*), environmental and genetic factors(*66*).

Our study found no group differences in blood biomarkers, including NFL and plasma pTau181. This lack of distinction may reflect the nonspecific nature of NFL, which indicates general neurodegeneration rather than AD-specific pathology(*31, 67*). Similarly, while pTau181 is associated with AD, its sensitivity for detecting early or preclinical stages is limited - emerging evidence suggests that other phosphorylated tau isoforms, such as pTau217, may offer greater diagnostic accuracy and specificity for AD-related pathology(*68*). Despite the absence of group differences, NFL predicted PI deficits, with associations observed for higher PI error, greater deviation from optimal velocity gain, and reporting noise. These associations align with NFL’s established link to systemic neurodegeneration and white matter pathology(*30, 69*), both critical for efficient neural communication(*70, 71*). Reduced white matter integrity, associated with elevated NFL, may amplify noise across neural networks, contributing to variability in velocity estimates and reporting accuracy. Furthermore, the NFL’s link to sensorimotor impairments, such as slower nerve conduction velocity and reduced sensory precision in diabetes(*72*), may further impact motor control and sensory integration, contributing to both higher reporting noise as well as precision of self-motion information that may affect internal velocity gain estimates. This interpretation is consistent with previous work by Stangl et al. (*17*) and Segen et al. (*19*), which suggested that path integration impairments in healthy aging arise primarily from degraded or noisy sensory inputs and computational processes, rather than from specific dysfunction in grid cell systems.

Together, our findings suggest that NFL reflects broader, age-related neuronal changes contributing to increased uncertainty in navigation, while memory leak consistently distinguished SCD from controls, likely indicating early entorhinal dysfunction. By disentangling these mechanisms, we enhance our ability to differentiate path integration impairments associated with normal aging from those linked to early Alzheimer’s disease— supporting the use of computational markers to identify individuals at greatest risk.

### Future directions and conclusion

Although the ultimate validation of PI as a predictive biomarker for Alzheimer’s disease will require longitudinal evidence, our current cross-sectional findings provide critical insights into early cognitive changes in individuals at risk. We recognize that the SCD population is heterogeneous and that premorbid variability in spatial navigation ability, along with factors such as age and sex, can influence individual PI performance. Moreover, some healthy older adults in our sample may also be on a trajectory toward cognitive decline, which longitudinal follow-up will help to clarify. Nevertheless, the group-level differences we observe— particularly the increased memory leak in SCD participants—demonstrate that even in the absence of overt cognitive impairment, subtle but systematic disruptions in navigational computations are already detectable. These findings highlight the sensitivity of self-motion-based path integration tasks for capturing early changes linked to Alzheimer’s disease risk and underscore the potential utility of computational modeling approaches for revealing latent cognitive vulnerabilities.

In summary, our findings highlight the potential of PI deficits—particularly increased memory leak—as early markers of Alzheimer’s disease risk in individuals with subjective cognitive decline. By decomposing PI errors using computational modeling, we revealed distinct mechanisms underlying navigational impairments that are not apparent with conventional cognitive assessments. These insights into early grid cell dysfunction and entorhinal cortex vulnerability can inform the development of targeted spatial navigation tasks for clinical use. Future work should determine whether PI-based measures can serve as sensitive endpoints for monitoring disease progression and evaluating the efficacy of disease-modifying interventions.

### Materials and Methods Participants

The study involved 104 participants, divided into two groups. The Control group consisted of 73 individuals (46 females), averaging 65.70 years old (SD = 5.80). The Subjective Cognitive Decline (SCD) group, referred by neurologists from an in-house memory clinic, included 31 participants (15 females), with an average age of 68.45 years (SD = 7.79). SCD classification was based on a comprehensive clinical interview, including self-reported cognitive concerns and informant feedback, with no objective cognitive impairment detected through neuropsychological testing using the CERAD-plus battery(*73*). All participants provided informed consent, and the study was approved by the Ethics Committee of the University of Magdeburg. Two subjects (1 SCD and 1 Control) scored below the Montreal Cognitive Assessment (MoCA(*74*) cutoff of 23(*75*) - indicating the presence of mild cognitive impairment - and were hence excluded from further analysis-resulting in the final sample of 102 participants (72 controls and 30 SCD. All subjects had normal or corrected-to-normal vision and were physically capable of standing for extended periods, a prerequisite for completing the PI task. We also obtained self-reported spatial abilities, measured by the 32-item DZNE Questionnaire on Spatial Orientation Skills (DFRO), and visuo-spatial working memory, measured by the Corsi block-tapping task(*37*), implemented using PsyToolkit platform(*76*). In addition to cognitive assessments, participants underwent functional gait analysis using four tasks from the Functional Gait Assessment(*24*), focusing on level surface walking, gait speed variations, narrow base support, and gait with eyes closed. Balance was assessed using eight brief 20-second tasks. However, due to scoring discrepancies among experimenters, these results were not included in the analysis.

### Plasma biomarker analysis

Blood samples for pTau181, NFL, NPTX2, and APOE genotyping analysis were obtained from 84 participants. The blood samples were analysed at the clinical research group, Bonn DZNE, using the SIMOA kit, whilst NPTX2 was analysed using the INNOTEST kit from Fujirebio.

We did not include NPTX2 in the final analysis because, although it is secreted by neurons and serves as a marker of synaptic integrity(*77*), NPTX2 is also produced in non-neuronal tissues such as the pancreas (pancreatic islets), pituitary gland, and adrenal medulla. This broader expression pattern raises concerns about the specificity of plasma NPTX2 as a reliable marker of synaptic integrity. For APOE genotyping, DNA was extracted from participants’ blood samples, analyzed to detect the APOE polymorphisms, and assigned two of the following alleles: ε2, ε3, or ε4. The APOE ε4 allele is a major risk factor of AD(*33*). We classified participants as ε4 carriers (ε3ε4, ε4ε4, and ε2ε4) or ε4 noncarriers (ε2ε2, ε2ε3, and ε3ε3).

### Immersive virtual reality path integration task

Participants engaged in a self-guided immersive virtual reality path integration task, performed in a virtual environment featuring an open field devoid of landmarks, with only a ground pebbly texture providing optic flow information. The self-guided nature of the task, where participants chose their preferred walking speed, offered the advantage of minimising experimenter biases and potential dual task costs associated with walking at a predefined speed. This setup also contrasts with other self-guided PI tasks, e.g., the apple game(*7*) or virtual reality-based triangle completion tasks (e.g.,(*9, 36*)) where external objects act as destination markers to guide participants, potentially enabling them to compute distances using static visual depth perception. This task required them to estimate the distance and direction to their starting point at two different points along each of eight unique sinuous paths - in the middle and at the end. These paths were designed with a variety of left and right turn combinations, ensuring each combination was repeated twice (Fig. S1). Examples include left followed by right turn, right followed by left turn, two consecutive left turns, and two consecutive right turns. The turn sizes varied between 40° and 140°, with the stipulation that the combined turn sizes in the same direction per path did not exceed 180°. This design, devoid of external guiding objects, ensured that distance estimation was based primarily on internal cues rather than visual distance estimation, thus providing a purer assessment of path integration abilities.

The task was developed using Unity software (19.4.0f1) and played through an HTC Vive Pro headset equipped with a wireless setup, enhancing the immersive experience. Each path segment, a portion of the path that contains a single turn in one direction, either leading from the start to the midpoint or from the midpoint to the end, spanned approximately 3 metres, varying with the curvature of the path (beeline distance of 2.7 metres). In about 10% of the trials, participants walked the entire path and provided responses only at the end, resulting in trials of shorter duration but covering the same distance. Each new trial commenced with participants walking towards an object, then facing the start of the path to memorise their position and heading orientation. They then followed a floating sphere to the first stopping point, where they provided both Angular Integration (AI) and Path Integration (PI) responses. After responding, participants were guided to continue the path by following the sphere until reaching the end, where AI and PI responses were again given. The order of AI and PI responses was counterbalanced among participants.

At each stopping point during the task, participants were asked to orient themselves towards their perceived starting position, using a virtual ruler projected on the ground to indicate the distance to this location. The line’s direction was controlled by the participant’s head movements, while its length was adjusted using the up and down keys on the HTC Vive controller.

Besides PI responses, we also obtained an AI response by asking participants to remember and recreate their initial heading orientation at each stopping point, achieved by physically rotating to their perceived initial heading and pressing the trigger on the HTC Vive controller. This task requires participants to both memorize their initial heading and update that heading throughout the trial as they navigate the curved path, thus capturing angular path integration rather than static orientation recall. This additional task, which was based on earlier work(*21*), aimed to assess participants’ ability to integrate heading changes (AI) without the confounding factor of distance integration, differing from standard approaches of decomposing the PI response into distance and angular error (see Segen et al.,(*19*) for further discussion).

### Experimental procedure

The study was conducted over two separate days, with sessions lasting three hours each. Participants initially engaged in six practice trials. The main trials were organised into blocks of 14, interspersed with mandatory short breaks. At the end of each block, participants undertook three additional distance estimation trials, requiring them to recall and then replicate specific distances - 1.4, 3.8, and 5.9 meters - using a virtual ruler, without physical movement. This task was included to investigate potential differences in visual distance estimation and response noise between the control and SCD groups.

A subset of the subjects in the control group performed PI tasks without the AI response, due to technical difficulties, we included these subjects in the analysis, as their PI error was similar to those who provided both the PI and AI responses (Fig. S11).

### Behavioral data analysis Outlier removal

A 2-step outlier removal procedure was applied. First, we removed trials where an accidental response was registered either due to technical issues or participants’ use of the controllers.

These trials were identified as follows: trials less than 2 seconds (lowest possible time), trials with distance responses less than.4 meters (minimum set distance), trials with identical distance to the random lengths of the line at the beginning of the response (within.01m threshold). We also removed all trials that had response times over 60 seconds (longer response times often accompanied by loss of connection, or interruptions due to clarifications from subjects about the task). The second step included removal of outliers based on PI task performance (PI and AI error) using Gaussian Mixture Modeling (GMM), to remove occasional trials where participants might have temporary lost concentration or got disoriented.

Specifically, outliers were identified using the densityMclust() function from the **mclust** R package, which fits a GMM to the empirical distribution of errors for each participant. Trials falling below the **5th percentile of the log-likelihood distribution** were flagged as outliers, as they represent data points with the lowest likelihood given the participant’s error profile.

Overall, this resulted in the exclusion of 10.77% of the data for PI error and 13.79% for AI error. We used the GMM-based outlier removal for distance estimation trials based on absolute distance error for each participant and each distance level, which resulted in the exclusion of 3.05% of the data

### Path integration metric calculation

The x and y coordinates of the presumed starting point according to the participant’s response were calculated by:

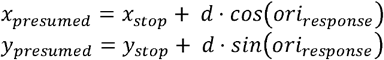

where d is the response distance, and *ori_response_* is the responded orientation. *x_origin_* and *y_origin_* are coordinates of the start point, *x_presumed_* and *y_presumed_* are the resulting coordinates of the presumed starting point. To determine the path integration error for a given stopping point, the Euclidean distance between the presumed starting point (according to the participant’s response at this respective stopping point) and the starting point was calculated

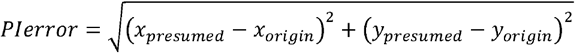

### Angular integration metric calculation

Angular integration error was calculated using the absolute difference between the initial heading orientation at the starting point (orientation indicated to participants using an arrow on the floor of the virtual environment) and the angular orientation response at each stopping point.

### Modelling analysis

#### Outlier removal

To model error sources, an additional outlier removal criterion was applied, excluding subjects with fewer than 50 valid PI trials after data pre-processing. This resulted in the removal of 10 subjects (7 controls and 3 SCD). Following parameter estimation, we further excluded subjects with negative velocity gain (α). This led to the exclusion of an additional 9 participants (3 Controls and 6 SCD). Examination of individual responses in this group revealed a common tendency to “fail” to turn during their PI response, contributing to the negative velocity gain. A detailed analysis of the error patterns and response profiles of these participants is provided in the supplementary materials.

Given that these 19 participants were excluded from the modelling analysis, we conducted a re-analysis of the behavioural data, also excluding these individuals, and present the results in the supplementary materials for comparison.

#### Internal estimate model

We used the distance model from Stangl et al.(*17*) where internal location estimates of the participants’ positions are modelled by a two-dimensional diffusion equation. Compared to Stangl et al.(*17*) where the path between two control points was approximated by a straight line, we interpolated the trajectories by a piecewise linear approximation. Bold-faced letters denote multi-dimensional vectors.

Let be a path of length parametrized by its length, i.e.,**x**(0) and **x**(*L*) correspond to the starting and the finishing point, respectively. Let 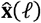 be the internal location estimate of the participant’s actual position x(*ℓ*) for 0≤*ℓ*≤ r. The distance model from Stangl et al.(*17*)

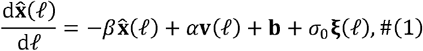

where:

- *β* is the location memory decay. If *β* = 0, the participant can incorporate the inputs on the right-hand side of Eq. (1) into the estimate of 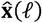 perfectly. If *β* > 0, the participant will slowly forget the previous inputs. Models of this type are known as “leaky integrators”.
- **V**(*ℓ*) = dx(*ℓ*)/d*ℓ* is the normalized velocity at x(*ℓ*). Since the path is parametrized by the distance, it follows that |v(*ℓ*)| =1 for all 0≤*ℓ*≤ *L*.
- *α* is the multiplicative velocity gain. The value a= 1 corresponds to the correct1<a evaluation of the contribution of v on the location estimate. The cases 0<a<1 and describe systematic underestimation and overestimation of the same effect, respectively.
- **b** i**s** the additive bias, i.e., the direction in which the internal estimate is being systematically shifted.
- *σ*_O_ is the accumulating noise (standard deviation). If *σ* _O_, the internal location estimate is not affected by the accumulating noise.
- ξ is two-dimensional normally distributed Gaussian noise uncorrelated in *ℓ*. Formally, the noise is a derivative of the two-dimensional Brownian motion.

We note that for *β*= 0, *σ*_0_ = 0, *α*=1 and b= u, the estimate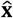 perfectly reflects the actual position.

### Segment reformulation

Assume that the path is split into segments marked by stopping points **s***_k_*, k= 0,1,2, …,*K*, so that **s_k_**=x (*ℓ_k_*) for some *ℓ_k_* ∈[0,*L*] with *ℓ*_0_=0 and *ℓ_K_* = *L.* Let *Δℓ_k_* = *ℓ_k_* -*ℓ*_*k*-1_, where *k*=1,2, …,*K* be the length of the *k*-th segment of the path. The internal estimate 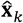 at the stopping point **s***_k_* can be recovered from the participant’s report of distance estimate 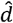 and the, estimate of angle 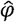 to the starting point x_start_ by

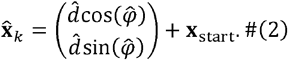

We set x_start_ = 0. Give-n the internal estimate 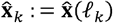 of location at the stopping point **s***_k_*, the internal estimate of 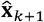 have a Gaussian distribution given by

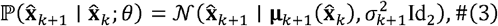

where (θ = β, α,**b**, σ_0_) are the model parameters, Id_2_ is the two-dimensional identity matrix, and the mean μ _*k*+1_ and the variance 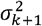 are defined by

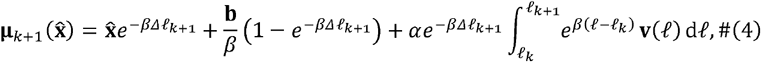

and

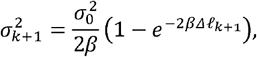

respectively (see supplemental material for complete derivation).

In Stangl et al.(*17*), the integral term in Eq. (4) is simplified by an additional assumption of a constant velocity along each segment, effectively approximating the trajectory of each segment by a straight line. In contrast, we have not imposed this additional assumption, which renders the integral analytically unsolvable in general. For our purposes, it was sufficient to employ a numerical method to approximate the integral with higher precision.

### Reporting noise

We consider reporting noise as a normal distribution with zero mean and variance 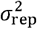 independent of ξ, reflecting the spread of the responses around the internally estimated location in Eq. (2). The reported internal location therefore satisfies:

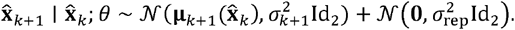

Thanks to the independence of ξ and the reporting noise, the density of the reported internal location simplifies to:

Following Weber’s law, we assume that the standard deviation of the reporting noise is proportional to the participants’ reported distance (at the end of the -th segment), i.e. :

where are the model parameters.

### Bayesian hierarchical model

We employed a Bayesian approach(*78*), MCMC sampling, to estimate the posterior distributions of the model parameters. The likelihood for a single path segment is given by Eq. (5). Consequently, the likelihood function for the whole path is:

For trials, let denote the vector of reports at the -th control point. The overall likelihood function is then defined by:

If represents the prior distribution over the parameters, the posterior distribution is:

We introduced two levels of hierarchy into each model parameter *ψ* : individual and group level, represented using the plate notation (Fig. 7). At the individual level, parameters from participants within the same group are assumed to follow the same prior distribution governed by the group-level parameters. Specifically, for a given parameter *ψ* associated with the *p* from group *g* (either Control or SCD) has a distribution 𝒟 with location *γ _ψ,g_* and scale*τ_ψ,g_*

**Fig. 7.**
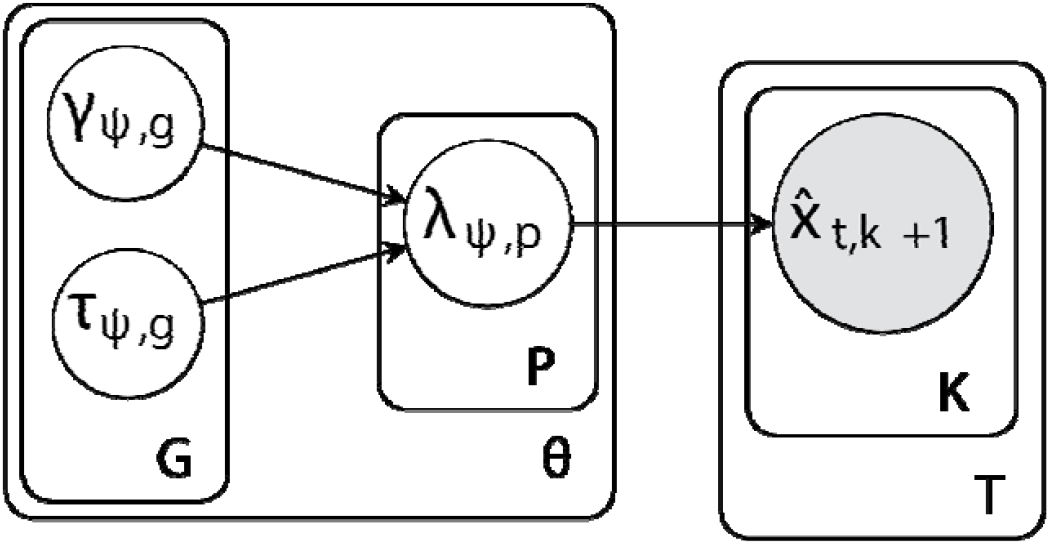
Graphical Representation of the Bayesian Hierarchical Model. The group-level hyper-parameters and, associated with group plate, govern the individual-level parameter, enclosed in the participant plate. Each participant undergoes multiple trials, represented by the outer trial plate, with each trial having multiple path segments captured by the inner plate. The observed data at segment in trial is influenced by the parameter. Here stands for any of five model parameters under parameter plate.

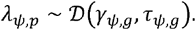

For accumulating noise *σ*_0_ and reporting noise is σ*_r_*, 𝒟 is Gaussian. For all other parameters 𝒟 is a Gaussian. The group-level hyper-parameters *γ _ψ,g_* and *τ_ψ,g_* have their own respective priors _ℋ 1_ and ℋ _2_:

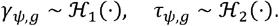

Details regarding the specific prior distribution of hyper-parameters, including their locations and scales, are provided in the supplemental material.

Since an analytical solution for the posterior distribution in Eq. (6) is not available, we used the No-U-Turn Sampler (NUTS) to generate posterior samples of the model parameters (*27*). The inference was conducted using NumPyro(*79*) with four independent MCMC chains, each run for 1000 warm-up iterations followed by 1000 sampling iterations. To assess model performance, we used leave-one-out expected log pointwise predictive density, elpd_loo_

### Statistics and reproducibility

#### PI and AI error analysis

For statistical quantification, all analyses were conducted in R. To examine the relationship between group status, stopping point, we used robust multiple linear regression with the MASS package in RStudio, as the Shapiro-Wilk test indicated non-normal residuals (p < 0.05). These models assessed associations of these factors with two primary outcomes: PI error (m) and AI error (°). Covariates included ‘sex’, ‘age’, and ‘MoCA’, and due to evidence suggesting sex-specific effects in AD pathology(*80*), a ‘sex by group’ interaction term was also added.

Continuous covariates were scaled and centred to normalize their range. We applied sum contrasts for binary factors such as group (control vs. SCD) and sex (male vs. female), and successive differences contrasts for stopping point, comparing intermediate versus final stopping points.

#### Blood and genetic biomarker analysis

To evaluate whether PI performance and key computational model parameters were related to biological and genetic markers of neuropathology (pTau 181, NFL and APOE status), we modelled PI error and parameters such as the absolute deviation from optimal velocity gain (1), beta, additive bias, accumulating noise, and reporting noise as dependent variables, influenced by standardised (scaled and centred) plasma biomarker concentrations. All models included a group interaction term and age as a covariate. Given the violation of normality, robust regression from the MASS package was employed to capture these relationships accurately. Sum contrasts were used for APOE status (carriers and noncarriers).

To evaluate the unique contribution of plasma NFL levels to specific error sources, partial R^2^ values were calculated. For each dependent variable (e.g., reporting noise, velocity gain), we compared the variance explained by full regression models including NFL with reduced models excluding NFL. Partial R^2^ was computed as the proportion of variance uniquely attributed to NFL, reflecting its specific predictive contribution to the model.

#### Group comparisons on demographic variables, blood biomarkers and movement characteristics

For simple group differences, Bayesian t-tests were conducted. Where variances were equal, we used ttestBF from the BayesFactor package in R; in cases of unequal variances, as in age, and gait, we modelled variance separately for each group using the brm function from the brms package. This method applied to demographic variables (age, MoCA, self-reported spatial abilities, visuo-spatial working memory, gait and number of completed trials) as well as group comparisons for blood biomarkers (pTau181, NFL) and movement metrics (head movements, angular and translational velocity, and head pitch).

For comparisons between first 10% and last 10% of trials on changes in PI performance and movement dynamics from early to late trials, we used linear regression analysis with sum contrasts for both group and trial period (first 10% and last 10%).

#### Modelling analysis

##### Individual-level

To examine differences for the individual (mean) level error sources, we used robust linear regressions from the MASS package to account for violations of the normality assumption in residuals. Separate models were fitted for each model parameter, with age included as a covariate. The parameters analyzed included memory leak (β), velocity gain (α), additive bias (‖**b**‖), accumulating noise 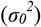, and reporting noise 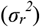. Sum contrasts were used for group were used (Control/SCD).

##### Group-level

For group-level analysis, we examined the posterior distributions of the model parameters to assess credible differences between groups. The analysis focused on the 95% Highest Density Interval (HDI), a key concept in Bayesian inference that indicates the range within which the most credible values of a parameter lie. Whether zero falls within this interval is crucial for interpreting the strength of evidence for an effect. If zero is excluded from the 95% HDI, it suggests statistically credible evidence of an effect, while inclusion of zero indicates the data do not rule out the possibility of no effect, reflecting uncertainty about the presence of a true difference. Additionally, we applied the Region of Practical Equivalence (ROPE)(*29*) to determine whether observed effects were practically negligible. The ROPE defines a range around the null value (often zero) within which differences are considered too small to be meaningful in practice. If most of the posterior distribution (e.g., 95% HDI) falls within the ROPE, the effect can be considered practically equivalent to the null value. We used ArviZ, NumPy, and Matplotlib to perform group-level analysis.

## Acknowledgements

We would like to thank the research assistants M. Schaumburg, F. Zeller, N. Behrenbruch, and C. Winter for their invaluable support in data collection, and to the medical staff, particularly F. Schulze and D. Hartmann, for their assistance with blood sample collection and S. Kuhs for biomarker analysis.

## Funding

This work was supported by:

Collaborative Research in Computational Neuroscience Grant (01GQ2106) of the German Ministry of Education and Research (BMBF)

Deutsche Forschungsgemeinschaft (DFG, Project-ID: 425899996 – SFB 1436) National Institutes of Health’s National Institute on Aging (5R01AG076198-02)

## Author contributions

Conceptualization: VS, TW, EN, ZT, MRK

Methodology: VS, TW, ZT, AS, JS, MRK, WG, MB

Investigation: VS, MRK

Visualization: VS, MRK

Supervision: TW, ZT

Writing—original draft: VS, TW

Writing—review & editing: VS, TW, EN, ZT, AS, JS, MRK, WG, MB

## Data and material availability

The summary and raw data used for the analyses, including results from the computational modeling, are available on Zenodo (https://doi.org/10.5281/zenodo.15574187). Code used for the computational modelling is published on Zenodo (https://doi.org/10.5281/zenodo.15532479) with latest updates available on GitHub (https://github.com/cogneuroai/Bayesian-hierarchical-model-for-PI). Code used to generate the plots presented in this manuscript is also available on Zenodo (https://doi.org/10.5281/zenodo.15574187).

