## Supplementary materials for "Path integration impairments reveal early cognitive changes in Subjective Cognitive Decline"

### Supplementary Text

**Derivation of formulas for $\boldsymbol{\mu}(\hat{\mathbf{x}}_{k+1})$ and $\sigma_{k+1}^2$:** We begin with a distance-based model for the internal location estimate (*17*):

$$\frac{\mathrm{d}\hat{\mathbf{x}}(\ell)}{\mathrm{d}\ell} = -\beta\hat{\mathbf{x}}(\ell) + \alpha\mathbf{v}(\ell) + \mathbf{b} + \sigma_0\boldsymbol{\xi}(\ell). \tag{S1}$$

Let $\mathbf{W}$ be a two-dimensional Brownian motion. We can write the model from Eq. (S1) as

$$\frac{\mathrm{d}\hat{\mathbf{x}}(\ell)}{\mathrm{d}\ell} + \beta\hat{\mathbf{x}}(\ell) = \alpha\mathbf{v}(\ell) + \mathbf{b} + \sigma_0\frac{\mathrm{d}\mathbf{W}(\ell)}{\mathrm{d}\ell}. \tag{S2}$$

By multiplying both sides of Eq. (S2) by an integrating factor $e^{\beta\ell}$ and using the product rule for derivatives with $\frac{\mathrm{d}}{\mathrm{d}\ell}e^{\beta\ell} = \beta e^{\beta\ell}$, we get

$$\frac{\mathrm{d}}{\mathrm{d}\ell}\left[\hat{\mathbf{x}}(\ell)e^{\beta\ell}\right] = e^{\beta\ell}\frac{\mathrm{d}\hat{\mathbf{x}}(\ell)}{\mathrm{d}\ell} + \beta e^{\beta\ell}\hat{\mathbf{x}}(\ell) = \alpha\mathbf{v}(\ell)e^{\beta\ell} + \mathbf{b}e^{\beta\ell} + \sigma_0 e^{\beta\ell}\frac{\mathrm{d}\mathbf{W}(\ell)}{\mathrm{d}\ell}.$$

Integrating both sides over $[\ell_k, \ell_{k+1}]$ gives

$$\begin{aligned}\hat{\mathbf{x}}_{k+1}e^{\beta\ell_{k+1}} - \hat{\mathbf{x}}_k e^{\beta\ell_k} &= \alpha\int_{\ell_k}^{\ell_{k+1}}\mathbf{v}(\ell)e^{\beta\ell}\,\mathrm{d}\ell + \mathbf{b}\int_{\ell_k}^{\ell_{k+1}}e^{\beta\ell}\,\mathrm{d}\ell + \sigma_0\int_{\ell_k}^{\ell_{k+1}}e^{\beta\ell}\,\mathrm{d}\mathbf{W}(\ell)\\ &= \alpha\int_{\ell_k}^{\ell_{k+1}}\mathbf{v}(\ell)e^{\beta\ell}\,\mathrm{d}\ell + \frac{\mathbf{b}}{\beta}\left(e^{\beta\ell_{k+1}} - e^{\beta\ell_k}\right) + \sigma_0\int_{\ell_k}^{\ell_{k+1}}e^{\beta\ell}\,\mathrm{d}\mathbf{W}(\ell).\end{aligned}$$

Using the notation $\Delta\ell_{k+1} := \ell_{k+1} - \ell_k$ and multiplying through by $e^{-\beta\ell_{k+1}}$ and inserting $1 = e^{\beta\ell_k}e^{-\beta\ell_k}$ in the integral terms, we obtain

$$\begin{aligned}\hat{\mathbf{x}}_{k+1} = \hat{\mathbf{x}}_k e^{-\beta\Delta\ell_{k+1}} + \frac{\mathbf{b}}{\beta}\left(1 - e^{-\beta\Delta\ell_{k+1}}\right) + \alpha e^{-\beta\Delta\ell_{k+1}}\int_{\ell_k}^{\ell_{k+1}}\mathbf{v}(\ell)e^{\beta(\ell-\ell_k)}\,\mathrm{d}\ell\\ + \sigma_0 e^{-\beta\Delta\ell_{k+1}}\int_{\ell_k}^{\ell_{k+1}}e^{\beta(\ell-\ell_k)}\,\mathrm{d}\mathbf{W}(\ell).\end{aligned}$$

From the properties of the stochastic integral, it follows that the distribution of $\hat{\mathbf{x}}_{k+1}$ given $\hat{\mathbf{x}}_k$ is Gaussian. Let us denote the mean and the variance matrix by $\boldsymbol{\mu}_{k+1}$ and $\mathbf{C} = (c_{ij})_{i,j=1}^2$ so that

$$\mathbb{P}\left(\hat{\mathbf{x}}_{k+1} \mid \hat{\mathbf{x}}_k\right) = \mathcal{N}\left(\boldsymbol{\mu}_{k+1}(\hat{\mathbf{x}}_k), \mathbf{C}\right).$$

Since the stochastic integral has zero mean value and the remaining terms are deterministic, the mean $\boldsymbol{\mu}_{k+1}(\hat{\mathbf{x}}_k)$ is given by the formula:

$$\begin{aligned}
\boldsymbol{\mu}_{k+1}(\hat{\mathbf{x}}_k) &= \mathbb{E}[\hat{\mathbf{x}}_{k+1}] \\
&= \mathbb{E}\bigg[\hat{\mathbf{x}}_k e^{-\beta\Delta\ell_{k+1}} + \frac{\mathbf{b}}{\beta}\left(1 - e^{-\beta\Delta\ell_{k+1}}\right) + \alpha e^{-\beta\Delta\ell_{k+1}} \int_{\ell_k}^{\ell_{k+1}} \mathbf{v}(\ell) e^{\beta(\ell-\ell_k)}\,\mathrm{d}\ell \\
&\qquad + \sigma_0 e^{-\beta\Delta\ell_{k+1}} \int_{\ell_k}^{\ell_{k+1}} e^{\beta(\ell-\ell_k)} \boldsymbol{\xi}(\ell)\,\mathrm{d}\mathbf{W}(\ell)\bigg] \\
&= \hat{\mathbf{x}}_k e^{-\beta\Delta\ell_{k+1}} + \frac{\mathbf{b}}{\beta}\left(1 - e^{-\beta\Delta\ell_{k+1}}\right) + \alpha e^{-\beta\Delta\ell_{k+1}} \int_{\ell_k}^{\ell_{k+1}} \mathbf{v}(\ell) e^{\beta(\ell-\ell_k)}\,\mathrm{d}\ell.
\end{aligned}$$

For the variance matrix, we need to employ the Itô isommetry which states that, given a one-dimensional Brownian motion $W$ and a suitable function $f$ and all $s \le t$, it holds

$$\mathbb{E}\left(\int_s^t f(r)\,\mathrm{d}W(r)\right)^2 = \int_s^t f^2(r)\,\mathrm{d}r. \tag{S3}$$

Hence, denoting the $i$-th component of a two-dimensional vector $\mathbf{y}$ by $\mathbf{y}_i$, we deduce

$$\begin{aligned}
c_{ij} &= \mathbb{E}\left[\left((\hat{\mathbf{x}}_{k+1})_i - (\boldsymbol{\mu}_{k+1})_i\right)\left((\hat{\mathbf{x}}_{k+1})_j - (\boldsymbol{\mu}_{k+1})_j\right)\right] \\
&= \sigma_0^2 e^{-2\beta\Delta\ell_{k+1}} \mathbb{E}\left[\left(\int_{\ell_k}^{\ell_{k+1}} e^{\beta(\ell-\ell_k)}\,\mathrm{d}\mathbf{W}_i(\ell)\right)\left(\int_{\ell_k}^{\ell_{k+1}} e^{\beta(\ell-\ell_k)}\,\mathrm{d}\mathbf{W}_j(\ell)\right)\right].
\end{aligned}$$

Since processes $\mathbf{W}_1$ and $\mathbf{W}_2$ are independent and the stochastic integrals have zero mean value, it holds

$$c_{12} = c_{21} = \sigma_0^2 e^{-2\beta\Delta\ell_{k+1}} \mathbb{E}\left(\int_{\ell_k}^{\ell_{k+1}} e^{\beta(\ell-\ell_k)}\,\mathrm{d}\mathbf{W}_1(\ell)\right) \mathbb{E}\left(\int_{\ell_k}^{\ell_{k+1}} e^{\beta(\ell-\ell_k)}\,\mathrm{d}\mathbf{W}_2(\ell)\right) = 0.$$

To compute $c_{11}$, we recall the Itô isometry from Eq. (S3) and observe

$$\begin{aligned}
c_{11} &= \sigma_0^2 e^{-2\beta\Delta\ell_{k+1}} \mathbb{E}\left(\int_{\ell_k}^{\ell_{k+1}} e^{\beta(\ell-\ell_k)}\,\mathrm{d}\mathbf{W}_1(\ell)\right)^2 \\
&= \sigma_0^2 e^{-2\beta\Delta\ell_{k+1}} \int_{\ell_k}^{\ell_{k+1}} e^{2\beta(\ell-\ell_k)}\,\mathrm{d}\ell \\
&= \frac{\sigma_0^2}{2\beta}\left(1 - e^{-2\beta\Delta\ell_{k+1}}\right).
\end{aligned}$$

The formula

$$c_{22} = \frac{\sigma_0^2}{2\beta}\left(1 - e^{-2\beta\Delta\ell_{k+1}}\right)$$

can be derived in exactly the same manner. Denoting $\sigma_{k+1}^2 := c_{11}$, we finally obtain

$$\mathbf{C} = \sigma_{k+1}^2\,\mathrm{Id}_2\,.$$

**Arc length parametrization:** Let us consider a path constructed by interpolating three control points $\mathbf{s}_0, \mathbf{s}_1, \mathbf{s}_2$ by a polynomial spline $\gamma : [0,2] \to \mathbb{R}^2$ satisfying $\gamma(k) = \mathbf{s}_k$ for $k = 0, 1, 2$. It is well-known that the length of the segment from $\gamma(0)$ to $\gamma(s)$ is

$$\mathrm{len}(s) = \int_0^s \left| \frac{\mathrm{d}\gamma(s)}{\mathrm{d}s} \right| \mathrm{d}s.$$

Let us denote $\ell_k = \mathrm{len}(k)$ for $k = 0, 1, 2$ so that $\ell_2$ is the total length of the path.

To numerically approximate the deterministic integral for the mean $\boldsymbol{\mu}_{k+1}(\hat{\mathbf{x}}_k)$, we need to be able to evaluate the velocity $\mathbf{v}(\ell)$ for any $0 \leq \ell \leq \ell_2$. This can be achieved by a parametrization map $\alpha : [0, \ell_2] \to [0, 2]$ such that the segment from $\gamma(0)$ to $\gamma(\alpha(\ell))$ has length $\ell$, in other words $\mathrm{len}(\alpha(\ell)) = \ell$. Informally, $\alpha$ maps length $\ell$ along the path to the index of control point. Since len is an increasing function, the inverse $\mathrm{len}^{-1}$ is well-defined and we may set

$$\alpha(\ell) = \mathrm{len}^{-1}(\ell).$$

Moreover, it can be shown that

$$\left| \frac{\mathrm{d}\gamma(\alpha(\ell))}{\mathrm{d}\ell} \right| = 1.$$

This identity reflects that the model in Eq. (S1) prescribes the change of the internal location estimate $\hat{\mathbf{x}}$ with respect to changes in distance $\ell$ along the path rather than with respect changes in to time. This corresponds to the distance model (*17*). Moreover, it can be shown that the derivative $\frac{\mathrm{d}\alpha(\ell)}{\mathrm{d}\ell}$ is the inverse of the magnitude of the velocity $\frac{\mathrm{d}\gamma}{\mathrm{d}s}$ at $s = \alpha(\ell)$.

In the code, we compute the functions len and $\alpha$ numerically. This allows us to approximate the deterministic integral by numerical integration.

**Hyper-parameters:** Locations and scales of the hyper-parameters:

$$\begin{aligned}
\gamma_{\alpha,g} &\sim \mathrm{Normal}(1, 10), & \tau_{\alpha,g} &\sim \mathrm{HalfCauchy}(5),\\
\gamma_{\beta,g} &\sim \mathrm{Normal}(0, 10), & \tau_{\beta,g} &\sim \mathrm{HalfCauchy}(5),\\
\gamma_{\mathbf{b},g} &\sim \mathrm{Normal}(0, 10), & \tau_{\mathbf{b},g} &\sim \mathrm{HalfCauchy}(5),\\
\gamma_{\sigma,g} &\sim \mathrm{Normal}_{+}(0, 1), & \tau_{\sigma,g} &\sim \mathrm{HalfCauchy}(5),\\
\gamma_{\sigma_r,g} &\sim \mathrm{Normal}_{+}(0, 1), & \tau_{\sigma_r,g} &\sim \mathrm{HalfCauchy}(5).
\end{aligned}$$

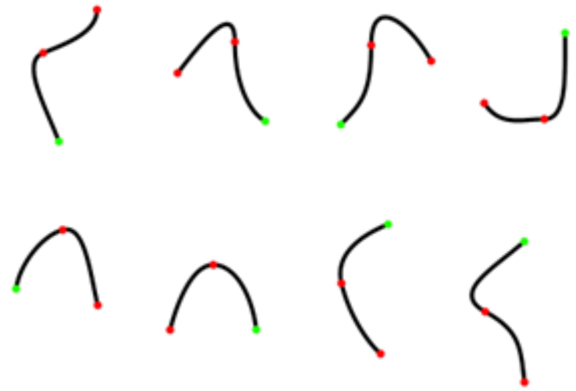

**Fig. S1. Path schematics used in the immersive path integration task.** Green point denotes start location, red points denote stopping points at which angular integration and path integration responses were given

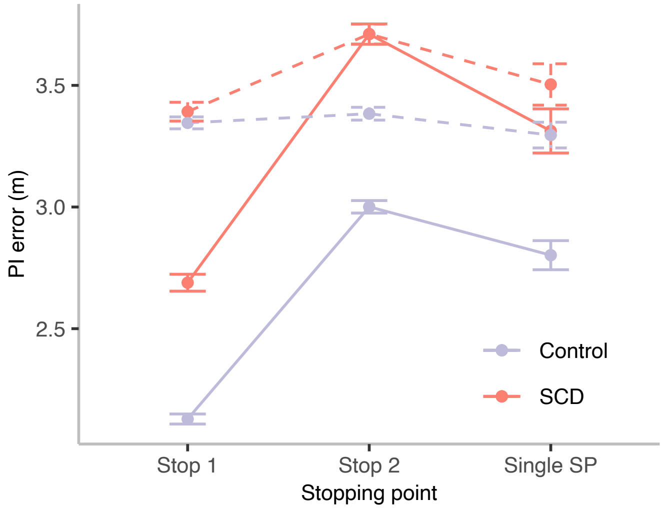

**Fig. S2. Path integration performance and comparison to chance level performance.** Path integration (PI) error as a function of stopping point, comparing performance between the Control and SCD groups. At Stop 1, both groups demonstrated performance significantly better than chance, with mean PI errors of 2.158 m for Controls (shuffled mean: 3.350, $p < 0.001$) and 2.692 m for SCD participants (shuffled mean: 3.384, $p = 0.004$). At Stop 2, Controls performed better than chance (Control: actual mean = 3.030, shuffled mean = 3.383, $p = 0.050$), while SCD were not significantly better than chance (SCD: actual mean = 3.723, shuffled mean = 3.730, $p = 0.979$). For the single stopping point condition, Controls performed significantly better than chance ($p = 0.012$) with a mean PI error of 2.854 m (shuffled mean: 3.311), whereas the SCD group did not (actual mean = 3.311, shuffled mean = 3.518; $p = 0.438$). Error bars represent the standard error of the mean.

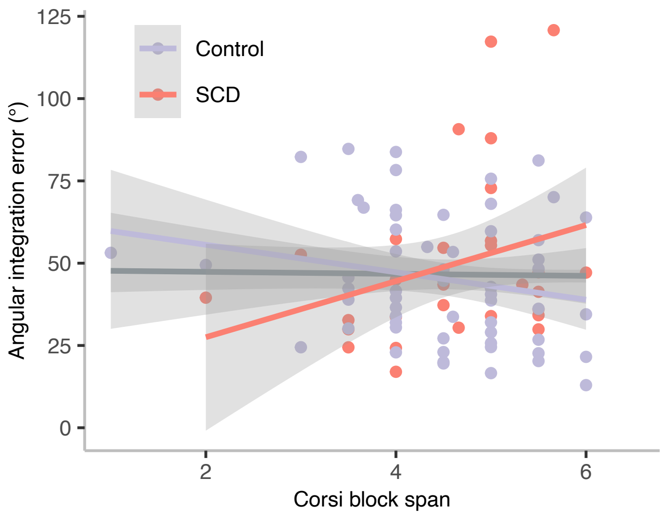

Fig. S3. **Link between AI error and Corsi block span, a standard measure of visuo-spatial working memory.** Each point represents an individual participant, with Control participants shown in purple and SCD participants in red. The colored lines indicate group-specific linear regression fits, with shaded areas representing 95% confidence intervals. The black line represents the overall linear fit across all participants. No significant correlation was observed between angular integration error and visuo-spatial working memory performance (Corsi span) in the full sample ($r = –0.013$, $p = 0.901$), nor within the control group ($r = –0.209$, $p = 0.108$) or the SCD group ($r = 0.302$, $p = 0.105$). These results suggest that working memory capacity, as measured by Corsi span, is unlikely to account for individual differences in AI error.

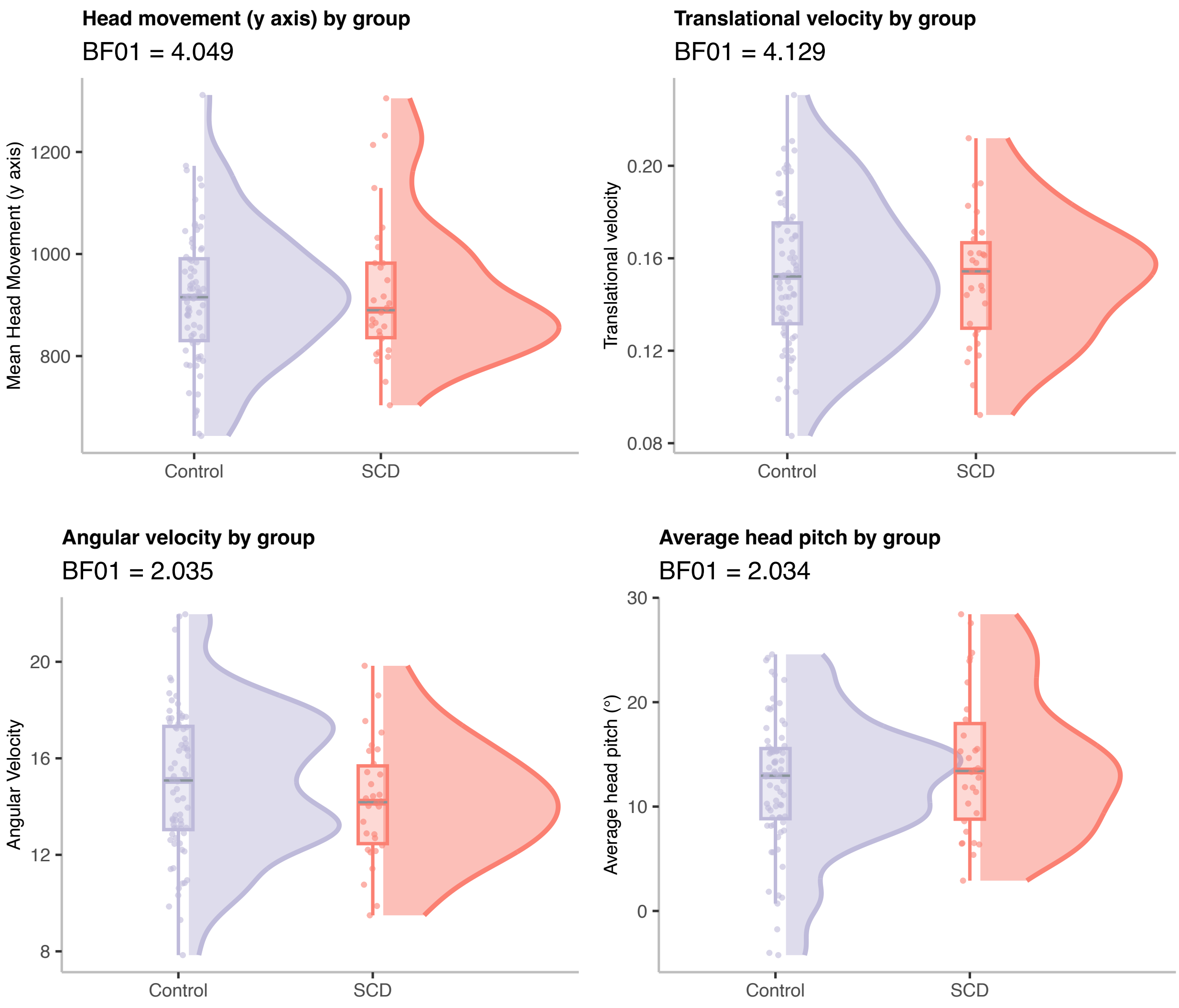

**Fig. S4. Comparison of Movement Dynamics Between Controls and SCD Participants**
Violin plots depicting the distributions of movement dynamics metrics across Control (blue) and SCD (red) groups together with overlaid boxplots representing the median and interquartile range (IQR), and individual data points. (Top left) Mean head movement along the y-axis ($BF_{01}$=4.049), (top right) translational velocity ($BF_{01}$=4.129), (bottom left) angular velocity ($BF_{01}$ =2.035), and (bottom right) average head pitch ($BF_{01}$ =2.034). Bayesian independent samples t-tests provided evidence in favor of the null hypothesis ($BF_{01}$ >1), indicating no significant group differences across these metrics. These findings suggest that differences in movement dynamics are unlikely to contribute to the observed differences in path integration errors between groups.

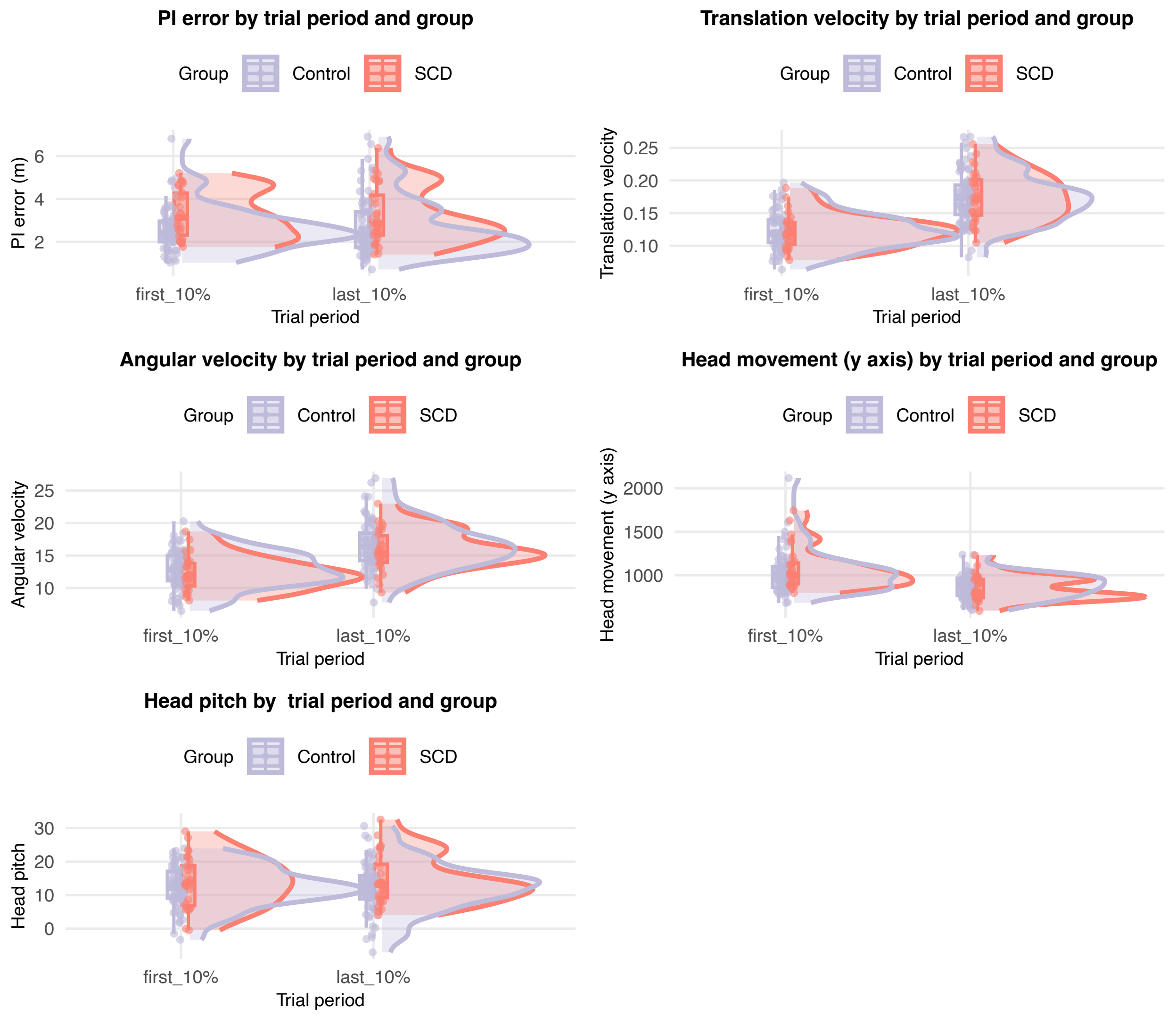

**Fig. S5. Path Integration Performance and Movement Dynamics Across Trial Periods and Groups.** Violin plots showing distributions of path integration (PI) error and movement dynamics metrics in SCD (red) and Control (blue) participants, separated by trial period (first 10% vs. last 10% of trials). (Top left) PI error remained consistent across trial periods for both groups, indicating no significant differences in learning dynamics or task adaptation. (Top right and middle left) Translational and angular velocities increased from early to late trials across both groups, with no significant group-by-trial interactions. (Middle right) Head movements decreased in later trials for both groups, potentially reflecting learning that distal cues were unavailable in the task environment. (Bottom) Head pitch showed no changes across trial periods or between groups. Together, these findings suggest that changes in movement dynamics or learning over the course of the task are unlikely to explain group differences in path integration performance. Each plot includes violin distributions showing data density, overlaid boxplots representing the median and interquartile range (IQR), and individual data points. Plots depict regression model results.

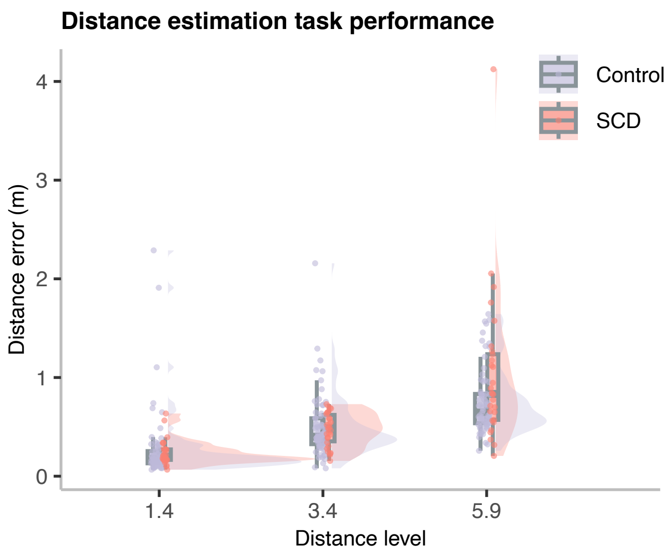

**Fig. S6. Distance Estimation Task Performance Across Groups**. Violin plots show distance estimation error for Control (gray) and SCD (red) participants across three target distances (1.4, 3.4, and 5.9 meters). Both groups exhibited a Weber-like pattern, with error increasing as target distance increased. No significant main effect of group was observed, though the SCD group showed a slightly larger increase in error at the longest distance, suggesting heightened distance-related uncertainty. Each plot includes violin distributions showing data density, overlaid boxplots representing the median and interquartile range (IQR), and individual data points. Full regression results are presented in the supplementary materials (Table S3).

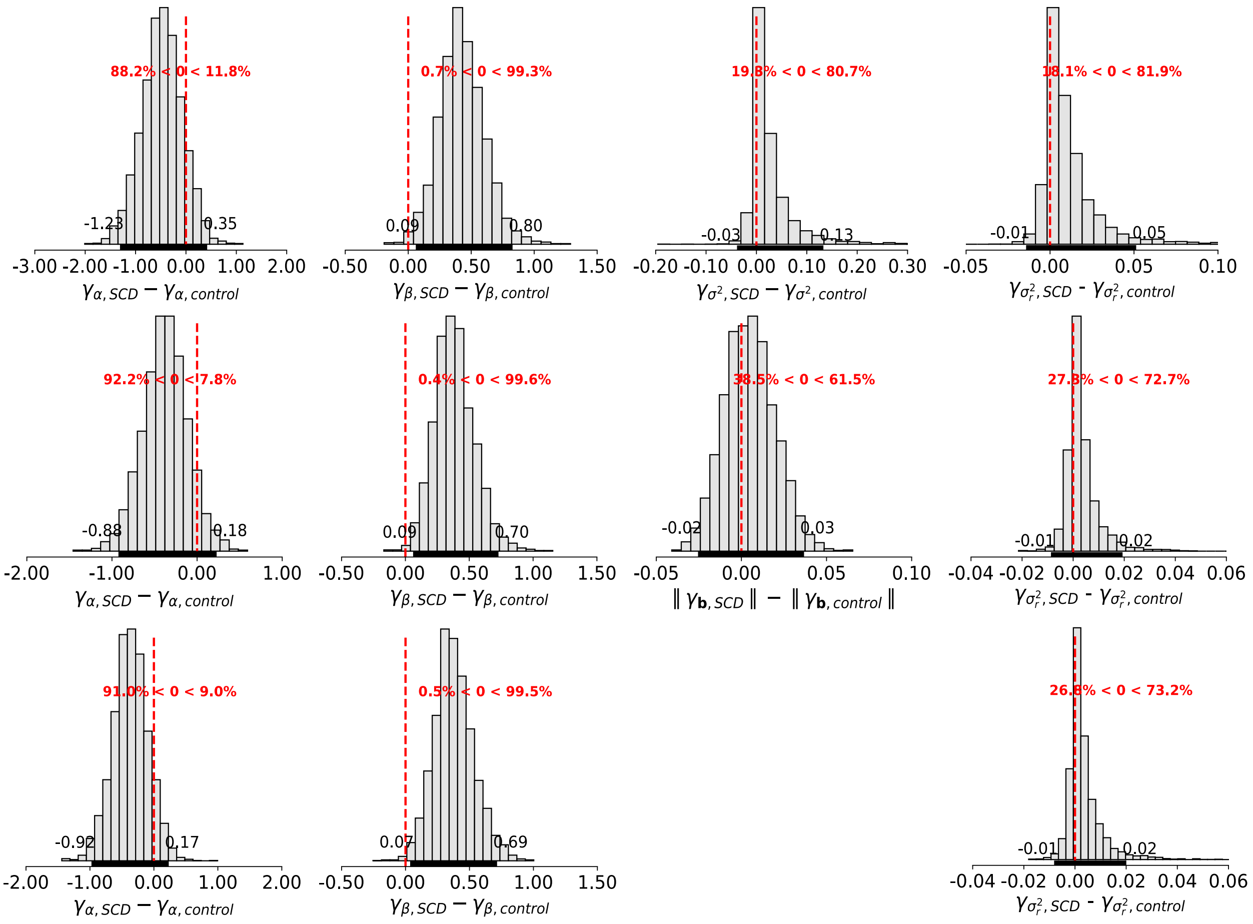

**Figure S7 Group-level posterior distributions of model parameter differences (SCD – Control) across three reduced versions of the computational model.** Each row represents a different reduced model: Top row: model with fixed additive bias (||b||), Middle row: model with fixed accumulating noise ($\sigma_0^2$), Bottom row: model with both additive bias (||b||) accumulating noise ($\sigma_0^2$) fixed. The histograms show posterior distributions of group differences for each parameter (velocity gain ($\alpha$), memory leak ($\beta$), reporting noise ($\sigma_r^2$) Horizontal black bars near the x-axis denote the 95% Highest Density Interval (HDI), while vertical dashed lines indicate zero. Percentages in red reflect the proportion of the posterior distribution on either side of zero, providing evidence for the likely direction of the group difference. Across all three reduced models, there was consistent and statistically credible evidence for higher memory leak ($\beta$), in individuals with SCD compared to controls, with the 95% HDI excluding zero and over 99% of the posterior distribution above zero. In contrast, the posterior distributions for velocity gain, additive bias, accumulating noise, and reporting noise showed negligible evidence for group differences, with 95% HDIs overlapping zero. These results replicate the findings from the full model and demonstrate that the memory leak effect is robust across alternative model specifications.

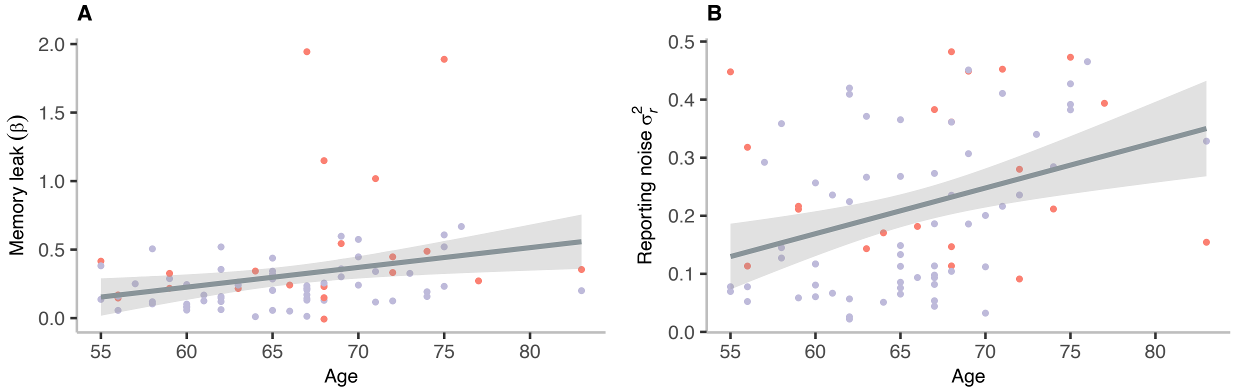

**Fig. S8. Scatter plots illustrating the relationship between age and model parameters across both groups**. Memory leak (β) and reporting noise ($\sigma_r^2$) increased significantly with age, independent of group. Shaded areas in scatterplots represent the 95% confidence interval.

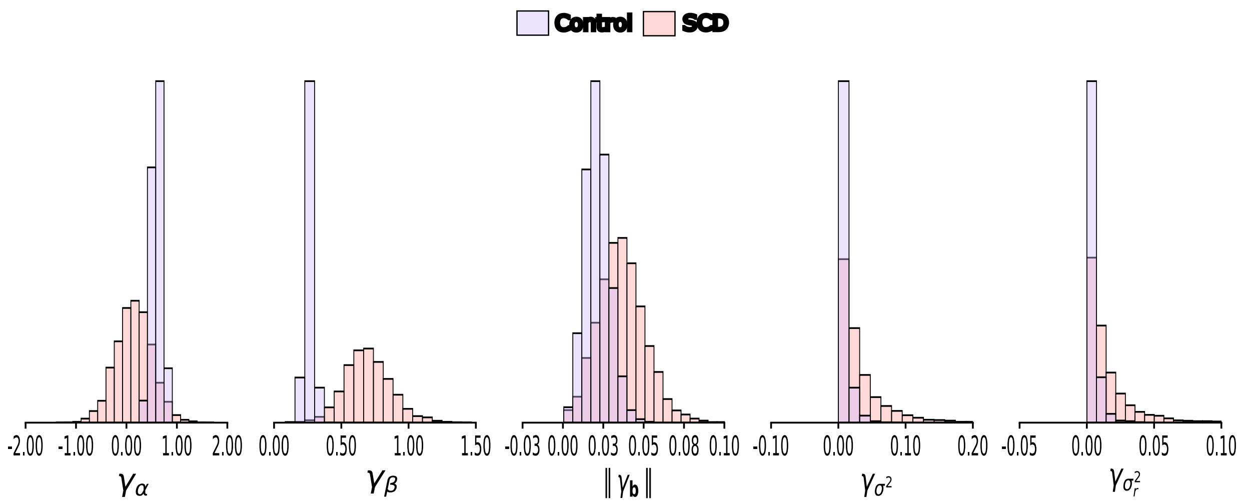

**Fig. S9. Posterior distributions of group-level parameters.** The marginal posterior distributions of the group-level parameters ($\gamma_\alpha$, $\gamma_\beta$, $\gamma_b$, $\gamma_{\sigma_0^2}$, and $\gamma_{\sigma_r^2}$) for SCD and Control.

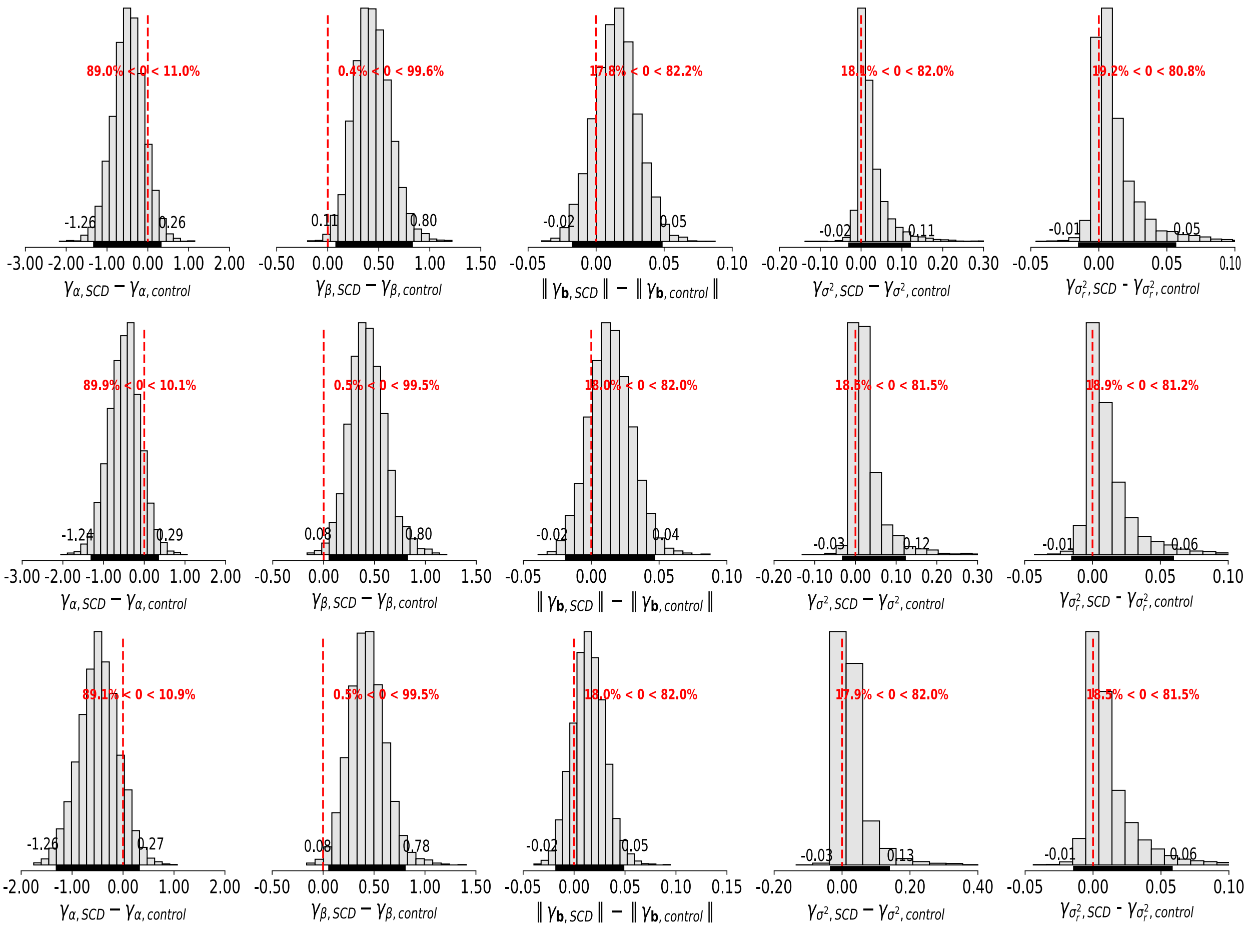

**Figure S10 Posterior differences for different scales of memory leak hyper-prior $\gamma_\beta$.** Each row corresponds to a different value for the prior scale on the standard deviation of the memory leak parameter: top row = 5, middle row = 10 (default used in the main manuscript), and bottom row = 15. The panels show posterior distributions of group differences for each model parameter. Results demonstrate that the group difference in memory leak ($\gamma_\beta$) remains consistent across different hyperparameter values, supporting the robustness of this effect to prior specification.

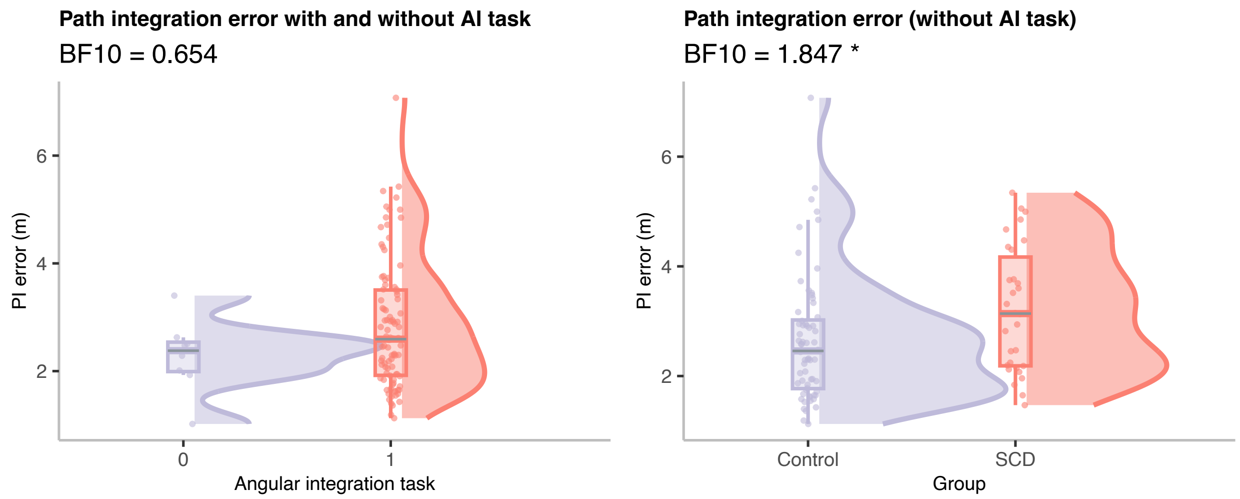

**Fig. S11. Path integration error comparisons with and without angular integration response** Eight subjects in our control group performed PI tasks without the AI response, to ensure that this has not affected our results (Left) Violin plot showing path integration (PI) errors for control participants who performed the PI task with and without the angular integration (AI) response ($BF_{10}$ =0.654), indicating no significant differences in PI error between these conditions. (Right) Violin plot showing PI errors for Control and SCD participants after excluding the eight control participants who did not perform the AI task. Significant group differences in PI errors were still observed ($BF_{10}$ =1.847), confirming that the inclusion of these participants did not influence the observed results. Each plot includes violin distributions to show data density, boxplots to display the median and interquartile range (IQR), and individual data points.

**Table S1. Robust multiple regression results for PI error (m)**

| | Full data set | | | | Subset used in modelling | | | |
|---|---|---|---|---|---|---|---|---|
| Predictors | Estimates | std. Error | Statistic | p | Estimates | std. Error | Statistic | p |
| (Intercept) | 2.783 | 0.063 | 44.101 | **<0.001** | 2.306 | 0.059 | 39.348 | **<0.001** |
| Group (*SCD*) | 0.250 | 0.064 | 3.899 | **<0.001** | 0.343 | 0.114 | 2.998 | **0.003** |
| Stop 1 vs Stop 2 | 0.491 | 0.088 | 5.586 | **<0.001** | 0.438 | 0.080 | 5.458 | **<0.001** |
| Stop 2 vs Single SP | 0.085 | 0.088 | 0.966 | 0.335 | 0.112 | 0.080 | 1.397 | 0.164 |
| Sex (*female*) | 0.284 | 0.066 | 4.311 | **<0.001** | 0.221 | 0.060 | 3.680 | **<0.001** |
| Age | 0.410 | 0.062 | 6.646 | **<0.001** | 0.291 | 0.052 | 5.633 | **<0.001** |
| MoCA | -0.019 | 0.060 | -0.320 | 0.749 | -0.068 | 0.051 | -1.330 | 0.185 |
| Group (*SCD*)× Stop 1 vs Stop 2 | 0.027 | 0.088 | 0.308 | 0.758 | 0.021 | 0.159 | 0.134 | 0.894 |
| Group (*SCD*)× Stop 2 vs Single SP | -0.047 | 0.088 | -0.538 | 0.591 | -0.104 | 0.159 | -0.657 | 0.512 |
| Group (*SCD*)× Sex (*female*) | 0.080 | 0.063 | 1.267 | 0.206 | 0.010 | 0.116 | 0.085 | 0.932 |
| Observations | 303 | | | | 246 | | | |

**Table S2. Robust multiple regression results for AI error (°)**

| | Full data set | | | | Subset used in modelling | | | |
|---|---|---|---|---|---|---|---|---|
| Predictors | Estimates | std. Error | Statistic | p | Estimates | std. Error | Statistic | p |
| (Intercept) | 42.836 | 1.352 | 31.672 | **<0.001** | 41.936 | 1.562 | 26.841 | **<0.001** |
| Group (*SCD*) | -0.190 | 1.367 | -0.139 | 0.889 | -3.206 | 2.962 | -1.083 | 0.280 |
| Stop 1 vs Stop 2 | 11.354 | 1.881 | 6.037 | **<0.001** | 10.243 | 2.153 | 4.758 | **<0.001** |
| Stop 2 vs Single SP | -2.181 | 1.881 | -1.160 | 0.247 | -2.701 | 2.153 | -1.255 | 0.211 |
| Sex (*female*) | 6.405 | 1.405 | 4.558 | **<0.001** | 8.221 | 1.599 | 5.143 | **<0.001** |
| Age | 3.407 | 1.347 | 2.529 | **0.012** | 3.161 | 1.370 | 2.308 | **0.022** |
| MoCA | -1.693 | 1.317 | -1.286 | 0.200 | -3.266 | 1.360 | -2.402 | **0.017** |
| Group (*SCD*)× Stop 1 vs Stop 2 | 0.634 | 1.881 | 0.337 | 0.736 | 2.493 | 4.122 | 0.605 | 0.546 |
| Group (*SCD*)× Stop 2 vs Single SP | 0.076 | 1.881 | 0.040 | 0.968 | -2.797 | 4.122 | -0.679 | 0.498 |
| Group (*SCD*)× Sex (*female*) | 0.500 | 1.348 | 0.371 | 0.711 | -4.654 | 3.015 | -1.544 | 0.124 |
| Observations | 285 | | | | 231 | | | |

**Table S3. Regression results for distance estimation task**

| | Distance error (m) | | | |
|---|---|---|---|---|
| Predictors | Estimates | std. Error | Statistic | p |
| (Intercept) | 0.090 | 0.231 | 0.390 | 0.697 |
| Group (*SCD*) | 0.027 | 0.024 | 1.113 | 0.267 |
| Distance (1.4 vs 3.4) | 0.288 | 0.033 | 8.722 | **<0.001** |
| Distance (3.4 to 5.9) | 0.351 | 0.033 | 10.553 | **<0.001** |
| Age | 0.007 | 0.003 | 1.978 | **0.049** |
| Group (*SCD*)× Distance (1.4 vs 3.4) | 0.056 | 0.033 | 1.706 | 0.089 |
| Group (*SCD*)× Distance (3.4 to 5.9) | 0.103 | 0.033 | 3.092 | **0.002** |
| Observations | 301 | | | |

**Table S4. Regression results for mean estimates for velocity gain bias, memory leak, additive bias, accumulating noise and reporting noise**

| | Velocity gain bias ($\alpha$) | | | |
|---|---|---|---|---|
| Predictors | Estimates | std. Error | Statistic | p |
| (Intercept) | 1.332 | 0.470 | 2.834 | **0.006** |
| Group (*SCD*) | -0.026 | 0.051 | -0.518 | 0.606 |
| age | -0.009 | 0.007 | -1.341 | 0.184 |

| | Memory leak ($\beta$) | | | |
|---|---|---|---|---|
| Predictors | Estimates | std. Error | Statistic | p |
| (Intercept) | -0.238 | 0.192 | -1.241 | 0.218 |
| Group (*SCD*) | 0.055 | 0.021 | 2.664 | **0.009** |
| age | 0.008 | 0.003 | 2.765 | **0.007** |

| | Additive bias ($\|\|\mathbf{b}\|\|$) | | | |
|---|---|---|---|---|
| Predictors | Estimates | std. Error | Statistic | p |
| (Intercept) | 0.026 | 0.024 | 1.075 | 0.285 |
| Group (*SCD*) | 0.001 | 0.003 | 0.570 | 0.571 |
| age | 0.000 | 0.000 | 0.376 | 0.708 |

| | Accumulating noise ($\sigma_0^2$) | | | |
|---|---|---|---|---|
| Predictors | Estimates | std. Error | Statistic | p |
| (Intercept) | -0.011 | 0.235 | -0.047 | 0.963 |
| Group (*SCD*) | 0.016 | 0.025 | 0.646 | 0.520 |
| age | 0.004 | 0.004 | 1.045 | 0.299 |

| | Reporting noise ($\sigma_r^2$) | | | |
|---|---|---|---|---|
| Predictors | Estimates | std. Error | Statistic | p |
| (Intercept) | -0.287 | 0.156 | -1.841 | 0.069 |
| Group (*SCD*) | 0.035 | 0.017 | 2.070 | **0.042** |
| age | 0.008 | 0.002 | 3.361 | **0.001** |
| Observations | 83 | | | |

**Table S5. Multiple regression results for PI error (pTau181, NFL and APOE4 genotyping as predictor variables)**

| | Full data set | | | | Subset used in modelling | | | |
|---|---|---|---|---|---|---|---|---|
| Predictors | Estimates | std. Error | Statistic | p | Estimates | std. Error | Statistic | p |
| (Intercept) | -0.422 | 1.299 | -0.325 | 0.746 | -0.107 | 1.192 | -0.090 | 0.929 |
| Group (*SCD*) | 0.429 | 0.502 | 0.856 | 0.395 | 0.411 | 0.415 | 0.991 | 0.326 |
| pTau181 | -0.150 | 0.421 | -0.356 | 0.723 | -0.101 | 0.376 | -0.270 | 0.788 |
| NFL | 1.080 | 0.366 | 2.954 | **0.004** | 0.761 | 0.321 | 2.371 | **0.021** |
| APOE4 Status (ε4 carriers) | 0.150 | 0.138 | 1.085 | 0.282 | 0.121 | 0.128 | 0.949 | 0.347 |
| Age | 0.037 | 0.021 | 1.735 | 0.087 | 0.031 | 0.020 | 1.608 | 0.114 |
| Group (*SCD*) x pTau181 | -0.339 | 0.435 | -0.780 | 0.438 | -0.082 | 0.388 | -0.212 | 0.833 |
| Group (*SCD*) x NFL | 0.188 | 0.330 | 0.570 | 0.570 | -0.066 | 0.297 | -0.223 | 0.824 |
| Group (*SCD*) x APOE4 Status (ε4 carriers) | 0.186 | 0.132 | 1.410 | 0.163 | 0.227 | 0.124 | 1.834 | 0.072 |
| | Observations 81 | | | | Observations 63 | | | |

**Table S6. Multiple regression results for individual parameter estimates derived from the computational model and (pTau181, NFL and APOE4 genotyping as predictor variables)**

| | Deviation from optimal velocity gain bias ($\alpha$) | | | |
|---|---|---|---|---|
| Predictors | Estimates | std. Error | Statistic | p |
| (Intercept) | 0.085 | 0.469 | 0.182 | 0.856 |
| Group (*SCD*) | 0.083 | 0.041 | 2.011 | **0.049** |
| pTau181 | -0.063 | 0.051 | -1.244 | 0.219 |
| NFL | 0.138 | 0.042 | 3.277 | **0.002** |
| APOE4 Status (ε4 carriers) | 0.083 | 0.046 | 1.807 | 0.076 |
| Age | 0.005 | 0.007 | 0.729 | 0.469 |
| Group (*SCD*) x pTau181 | 0.016 | 0.052 | 0.297 | 0.768 |
| Group (*SCD*) x NFL | 0.025 | 0.039 | 0.655 | 0.515 |
| Group (*SCD*) x APOE4 Status (ε4 carriers) | 0.094 | 0.044 | 2.113 | **0.039** |
| | Memory leak ($\beta$) | | | |
| Predictors | Estimates | std. Error | Statistic | p |
| (Intercept) | -0.081 | 0.264 | -0.307 | 0.760 |
| Group (*SCD*) | 0.078 | 0.023 | 3.365 | **0.001** |
| pTau181 | 0.004 | 0.029 | 0.148 | 0.883 |
| NFL | 0.043 | 0.024 | 1.793 | 0.079 |
| APOE4 Status (ε4 carriers) | -0.000 | 0.026 | -0.019 | 0.985 |
| Age | 0.006 | 0.004 | 1.404 | 0.166 |
| Group (*SCD*) x pTau181 | 0.033 | 0.029 | 1.107 | 0.273 |
| Group (*SCD*) x NFL | 0.005 | 0.022 | 0.214 | 0.831 |
| Group (*SCD*) x APOE4 Status (ε4 carriers) | 0.013 | 0.025 | 0.515 | 0.608 |

| | Additive bias ($||\mathbf{b}||$) |
|---|---|---|---|---|
| Predictors | Estimates | std. Error | Statistic | p |
| (Intercept) | 0.035 | 0.029 | 1.228 | 0.225 |
| Group (*SCD*) | -0.001 | 0.010 | -0.122 | 0.903 |
| pTau181 | 0.010 | 0.009 | 1.062 | 0.293 |
| NFL | 0.005 | 0.008 | 0.685 | 0.496 |
| APOE4 Status (ε4 carriers) | -0.005 | 0.003 | -1.721 | 0.091 |
| Age | -0.000 | 0.000 | -0.471 | 0.639 |
| Group (*SCD*) x pTau181 | 0.007 | 0.009 | 0.740 | 0.462 |
| Group (*SCD*) x NFL | -0.004 | 0.007 | -0.522 | 0.604 |
| Group (*SCD*) x APOE4 Status (ε4 carriers) | -0.000 | 0.003 | -0.133 | 0.895 |
| | Accumulating noise ($\sigma_0^2$) | | | |
| Predictors | Estimates | std. Error | Statistic | p |
| (Intercept) | -0.089 | 0.312 | -0.285 | 0.776 |
| Group (*SCD*) | -0.085 | 0.109 | -0.784 | 0.437 |
| pTau181 | 0.144 | 0.098 | 1.470 | 0.147 |
| NFL | -0.013 | 0.084 | -0.150 | 0.881 |
| APOE4 Status (ε4 carriers) | -0.055 | 0.033 | -1.647 | 0.105 |
| Age | 0.003 | 0.005 | 0.536 | 0.594 |
| Group (*SCD*) x pTau181 | 0.118 | 0.102 | 1.162 | 0.250 |
| Group (*SCD*) x NFL | -0.002 | 0.078 | -0.030 | 0.976 |
| Group (*SCD*) x APOE4 Status (ε4 carriers) | -0.025 | 0.032 | -0.788 | 0.434 |
| | Reporting noise ($\sigma_r^2$) | | | |
| Predictors | Estimates | std. Error | Statistic | p |

| | | | | |
|---|---|---|---|---|
| (Intercept) | -0.118 | 0.208 | -0.569 | 0.572 |
| Group (*SCD*) | 0.052 | 0.018 | 2.863 | **0.006** |
| pTau181 | -0.030 | 0.022 | -1.350 | 0.183 |
| NFL | 0.053 | 0.019 | 2.869 | **0.006** |
| APOE4 Status (ε4 carriers) | 0.033 | 0.020 | 1.630 | 0.109 |
| Age | 0.005 | 0.003 | 1.744 | 0.087 |
| Group (*SCD*) x pTau181 | -0.011 | 0.023 | -0.493 | 0.624 |
| Group (*SCD*) x NFL | -0.012 | 0.017 | -0.716 | 0.477 |
| Group (*SCD*) x APOE4 Status (ε4 carriers) | 0.034 | 0.020 | 1.723 | 0.091 |
| Observations | | | | 63 |

**Movie S1. Example of the Immersive Virtual Reality Path Integration Task.**
This video showcases two example trials from the immersive path integration task, presented in a first-person view. The first trial includes two stopping points, while the second features a single stopping point. In both trials, participants first complete the path integration response, followed by the angular integration response.
